# Transcriptional down-regulation of metabolic genes by Gdown1 ablation induces quiescent cell re-entry into the cell cycle

**DOI:** 10.1101/2020.03.01.971945

**Authors:** Miki Jishage, Keiichi Ito, Chi-Shuen Chu, Xiaoling Wang, Masashi Yamaji, Robert G. Roeder

## Abstract

Liver regeneration and metabolism are highly interconnected. Here, we show that hepatocyte-specific ablation of RNA polymerase II (Pol II)-associated Gdown1 leads to down-regulation of highly expressed genes involved in plasma protein synthesis and metabolism, a concomitant cell cycle re-entry associated with induction of cell cycle-related genes (including *cyclin D1*). and up-regulation of *p21* through activation of p53 signaling. In the absence of p53, Gdown1-deficient hepatocytes show a severe dysregulation of cell cycle progression, with incomplete mitoses, and a pre-malignant-like transformation. Mechanistically, Gdown1 is associated with elongating Pol II on the highly expressed genes and its ablation leads to reduced Pol II recruitment to these genes, suggesting that Pol II redistribution may facilitate hepatocyte re-entry into the cell cycle. These results establish an important physiological function for a Pol II regulatory factor (Gdown1) in the maintenance of normal liver cell transcription through constraints on cell cycle re-entry of quiescent hepatocytes.

## Introduction

Hepatocytes are major players in carrying out liver functions, such as nutrient metabolism and synthesis of plasma proteins. Although they rarely divide, hepatocytes re-enter the cell cycle upon liver injury or loss to restore liver mass (Michalopoulos, 2017). For decades, the molecular mechanism of liver regeneration has been intensively studied to identify factors that regulate the regeneration, and these studies have unveiled signaling pathways associated with cytokines, growth factors, and transcription factors (Michalopoulos, 2007). Although no studies have identified a single factor whose deletion abolishes liver regeneration (Michalopoulos, 2014), a recent study showed that the combined elimination of receptor tyrosine kinases, MET and epidermal growth factor receptor (EGFR) abolishes liver regeneration (Paranjpe et al., 2016). The study also showed that the elimination of MET and EGFR down-regulates genes involved in metabolic activities. In this regard, once quiescent cells commit to cell cycle entry, cellular metabolic activities must be changed in order to produce the components needed for cell doubling and cell survival. During liver regeneration, metabolic remodeling occurs along with cell division; and conversely, metabolic deficiency can impair regeneration (Caldez et al., 2018; Huang and Rudnick, 2014). Therefore, liver regeneration and metabolism are interconnected.

Normal hepatocytes exhibit a liver-specific pattern of gene expression that is altered in response to conditions leading to liver regeneration and altered cell metabolism, and the gene expression programs are regulated at least in part at the level of transcription (Hirota and Fukamizu, 2010; Kurinna and Barton, 2011). The transcription of protein coding genes, as well as some genes producing snRNAs and microRNAs, is mediated by RNA polymerase II (Pol II) in association with a group of general initiation factors (TFIIA, TFIIB, TFIID, TFIIE, TFIIF and TFIIH) that (with Pol II) form a preinitiation complex (PIC) at the promoter and by several elongation factors that facilitate promoter clearance and productive elongation (Roeder, 2019). The gene-selective formation and function of preinitiation complexes is in turn regulated by gene- and cell-specific enhancer-binding transcription factors that act through interactions with diverse transcriptional coactivators and corepressors. Foremost among the transcriptional coactivators is the large 30-subunit Mediator that is recruited by enhancer-bound transcription factors and, through Pol II interactions, facilitates enhancer-promoter interactions leading to PIC formation and function (Malik and Roeder, 2010).

Pol II(G) is a Pol II variant that contains Gdown1. Gdown1 was originally recognized as one of the multiple polypeptides encoded in the GRINL1A region of the human genome (Roginski et al., 2004). It later was identified biochemically as a stochiometric and tightly associated subunit (defined as the Pol II subunit, *Polr2m*) of a fraction of total Pol II purified from liver (Hu et al., 2006). Initial in vitro transcription assays reconstituted with purified initiation factors showed that Gdown1 inhibits transcription initiation but that Mediator can reverse this inhibition. Further biochemical and structural studies showed that Gdown1 blocks initiation by preventing interactions of initiation factors TFIIF and TFIIB with Pol II (Jishage et al., 2012; Jishage et al., 2018), although the mechanism by which Mediator reverses Gdown1-mediated repression remains unclear. These results indicate a new mechanism of transcriptional regulation in which Gdown1 directly interacts with Pol II to restrict potentially inappropriate Pol II recruitment to promoter regions. However, there is little information regarding the biological functions of Pol II(G), and how the clearly evident in vitro repressive function of Gdown1 relates to transcriptional regulation in vivo.

In this study, prompted in part by the original discovery of an apparently higher population of Pol II(G) relative to Pol II in porcine liver (Hu et al., 2006), the inhibitory nature of Gdown1, and the quiescent state of hepatocytes, we analyzed the function of Gdown1 in liver by a genetic analysis. We show that Gdown1 ablation in hepatocytes causes a surprising down-regulation of highly expressed genes involved in metabolic pathways and synthesis of plasma proteins in the liver, and that this triggers hepatocyte to re-enter into the cell cycle. We also show that the joint ablation of Gdown1 and p53 leads to dysregulated cell cycle progression, thereby implicating metabolic reprogramming in the molecular mechanism of malignant transformation.

## Results

### Gdown1 is essential for mouse early embryonic development

As an initial approach to investigate the biological role of Gdown1, we generated mice (*Gdown1^f/f^*; designated FF mice) carrying floxed exons in the *Gdown1* locus, such that Cre-recombinase expression excises exons encoding domains critical for the transcriptional inhibitory activity of Gdown1 (Supplemental Fig. S1A) (Jishage et al., 2018). First, we examined the effect of *Gdown1* ablation in mouse embryo development. *Gdown1^f/-^* mice appeared normal and healthy. However, no *Gdown1****^-/-^*** mice from intercrossing *Gdown1^f/-^* mice were obtained at postnatal day 21 (Supplemental Fig. S1B), indicating that *Gdown1* knockout (KO) mice are embryonic lethal. Further analyses revealed that the number of *Gdown1****^-/-^*** embryos began to decrease at embryonic day 3.5 (E3.5) and that no nullizygous embryos were evident at E10.5 (Supplemental Fig. S1B). Since *Gdown1* KO embryos were observed at E3.5, we generated embryonic stem cell (ESC) lines from *Gdown1****^f/f^*** (FF) mice carrying a tamoxifen-inducible *Cre-ERT2* transgene in order to examine the impact of Gdown1 loss on ESCs. In a defined culture condition, more than 60 ESC clones were screened by a single colony culture to identify *Gdown1* KO ESCs. However, the clones that survived after tamoxifen treatment were all heterozygous, suggesting that *Gdown1* KO ESCs may die very quickly. Taken together, these results show that Gdown1 is critical for mouse early embryonic development.

### Loss of Gdown1 activates p53 signaling pathway

Because of the embryonic lethality, we next chose mouse liver to investigate the biological role of Gdown1. While ESCs divide every 24 hours, normal hepatocytes are quiescent and do not divide frequently (Fausto et al., 2006; Michalopoulos, 2007; Taub, 2004). We generated *Gdown1^flox/flox^* mice carrying an *albumin(Alb)-Cre* transgene (*Gdown1^f/f;Alb-Cre^* mice; designated KO mice), and the hepatocyte-specific deletion of the *Gdown1* targeted allele was confirmed (Supplemental Fig. S1C). At 8 weeks, Gdown1 protein expression in *Gdown1* KO liver was barely detectable compared to expression of the Pol II RPB3 subunit (Supplemental Fig. S1D). In contrast to the observed lethal phenotype in *Gdown1* KO embryos, KO mice displayed whole body and liver weights comparable to those of control (FF) mice (Supplemental Fig. S1E). However, liver function tests showed abnormal liver metabolic activities, such as elevated serum levels of alkaline phosphatase (ALP) and lowered serum levels of triglycerides (TRIG) (Fig. 1A), that suggested functional defects in KO liver. Histological analyses of KO liver showed further hepatocyte abnormalities such as large cells with enlarged nuclei (H&E in Fig. 1B). Also, significant numbers of KO hepatocytes were Ki67-positive, which indicates that these cells had re-entered the cell cycle. While cell cycle re-entry was observed, a TUNEL assay detected a few apoptotic hepatocytes in *Gdown1* KO liver (Fig. 1B). Although obvious necrotic lesions were not detected, the presence of apoptotic cells may explain the proliferation of SMA-positive cells associated with collagen deposition (Fig. 1B), whose emergence is often triggered by hepatic injury (Yin et al., 2013).

**Figure 1.**
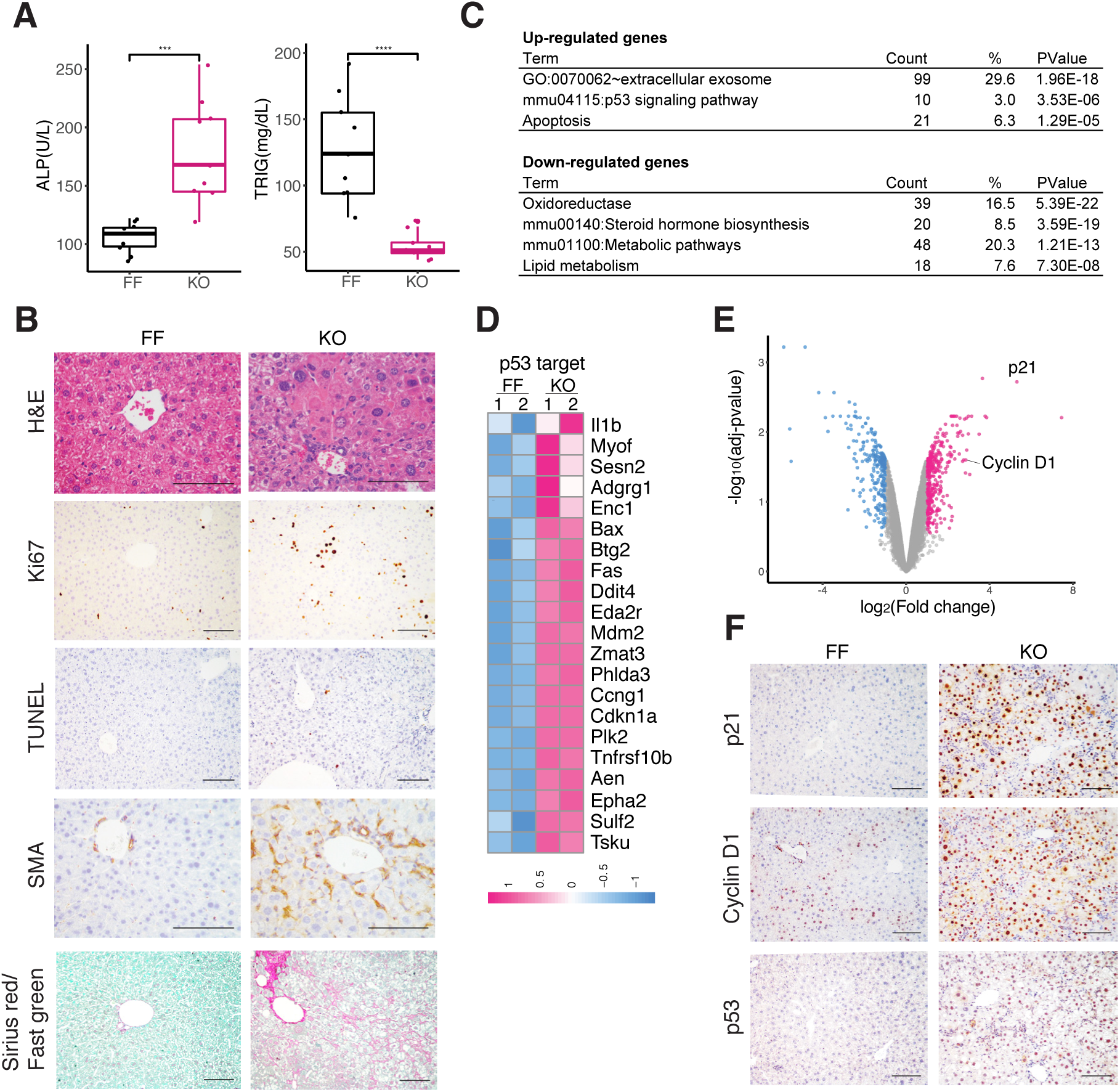
Loss of Gdown1 activates p53 signaling pathway. (A) Serum levels of alkaline phosphatase (ALP) and triglycerides (TRIG) in control (FF) and *Gdown1* liver KO mice (KO). \*\*\**P* < 0.001, \*\*\*\**P* < 0.0001 in unpaired two-tailed t-test. (B) Representative liver histology in control (FF) and *Gdown1* liver KO mice (KO). (C) Gene ontology analysis for differentially expressed genes in *Gdown1* KO liver. (D) A heatmap for p53 target genes that are up-regulated in *Gdown1* KO liver. (E) Volcano plot showing differentially expressed genes in *Gdown*1 KO liver. Deep pink or light blue color indicates significantly up- or down-regulated genes, respectively (Fold change > 2, p value < 0.05). (F) Representative liver histology in FF and *Gdown*1 KO mice. All scale bars represent 100 μm.

To understand the injury-like reactions, the transcriptome in *Gdown1* KO liver was analyzed by microarrays. We identified 338 up-regulated genes and 246 down-regulated genes (fold change > 2, p value < 0.05). GO analysis revealed that the up-regulated genes are significantly enriched in extracellular exosome, p53 signaling and apoptosis pathways, whereas the down-regulated genes are highly involved in metabolic pathways that include xenobiotic metabolism and lipid metabolism (Fig. 1C). Among genes that are categorized as “extracellular exosome” – and which also include genes localized in the plasma membrane, the Golgi apparatus, and the endoplasmic reticulum -- we found several genes that are normally expressed in cholangiocytes (KRT19-positive cells) and that include *Spp1*, *Epcam*, and *Prom1* (Supplemental Fig. S1F, G). Histological analyses confirmed the proliferation of cholangiocytes (Supplemental Fig. S1H), which is generally associated with biliary injury or exposure to alcohol, toxins or drugs (Alvaro et al., 2007). Although GO analysis identified only 10 genes involved in p53 signaling pathways (Fig. 1C), we found more genes that are direct targets of p53 (Fischer, 2017) -- including pro-apoptotic genes such as *Bax*, *Fas*, and *Tnfrsf10b* (Fig. 1D). In particular, *Cdkn1a/p21*, which encodes the potent cell cycle inhibitor p21, was remarkably up-regulated (Fig. 1E). Also, beyond genes in p53 signaling pathways, we found that *cyclin D1* was also significantly up-regulated (Fig. 1E). Histological analyses further confirmed expression at the protein level of cyclinD1, as well as p21 and p53, all of which were highly and uniformly expressed in KO liver (Fig. 1F). These results establish that *Gdown1* KO in hepatocytes causes injury-like reactions that include cell death, cell cycle re-entry, and activation of p53 signaling pathways in the liver.

### *Gdown1* KO hepatocytes re-enter into the cell cycle

To investigate how the *Gdown1* KO elicits injury-like reactions, we monitored expression of p21 and cyclin D1 proteins from 4-6 week (W) old KO livers. Cyclin D1 and p21 were both detected at 5W, when Gdown1 expression was nearly undetectable (Fig. 2A, lane 4), and further increased at 6W (lane 6), indicating that activation of a p53 signaling pathway was initiated around 5W of age. The number of Ki67-positive cells was elevated in KO liver at 5W, but followed by a decline at 6W (Fig. 2B). The decreased number of Ki67-positive cells at 6W was associated with *p21* induction accompanied by the expression of other p53 direct target genes that included *Phlda3*, *Zmat3*, and *Eda2r* (Supplemental Fig. S2), suggesting that the induced cell cycle re-entry and the subsequent progression was prevented through *p21* induction. Consistent with the observation of an increased number of Ki67-positive cells at 5W, serine 10-phosphorylated histone H3 also was detected at this stage (Fig. 2C), indicating that the cell cycle had progressed into the mitotic phase in some cells. Increases in both cyclin A2 and cyclin B2 RNA expression levels at 5W further confirmed that the hepatocytes had re-entered the cell cycle at this stage (Fig. 2D). However, no obvious necrosis or SMA-positive myofibroblasts or apoptotic hepatocytes were detected at 5W (Fig. 2E), suggesting that hepatic injury is unlikely to be the major cause of the cell cycle re-entry. Interestingly, expression of both cyclin D1 mRNA and protein was observed at 6W (Fig. 2F), even though the cell cycle appeared to be arrested (Fig. 2B).

**Figure 2.**
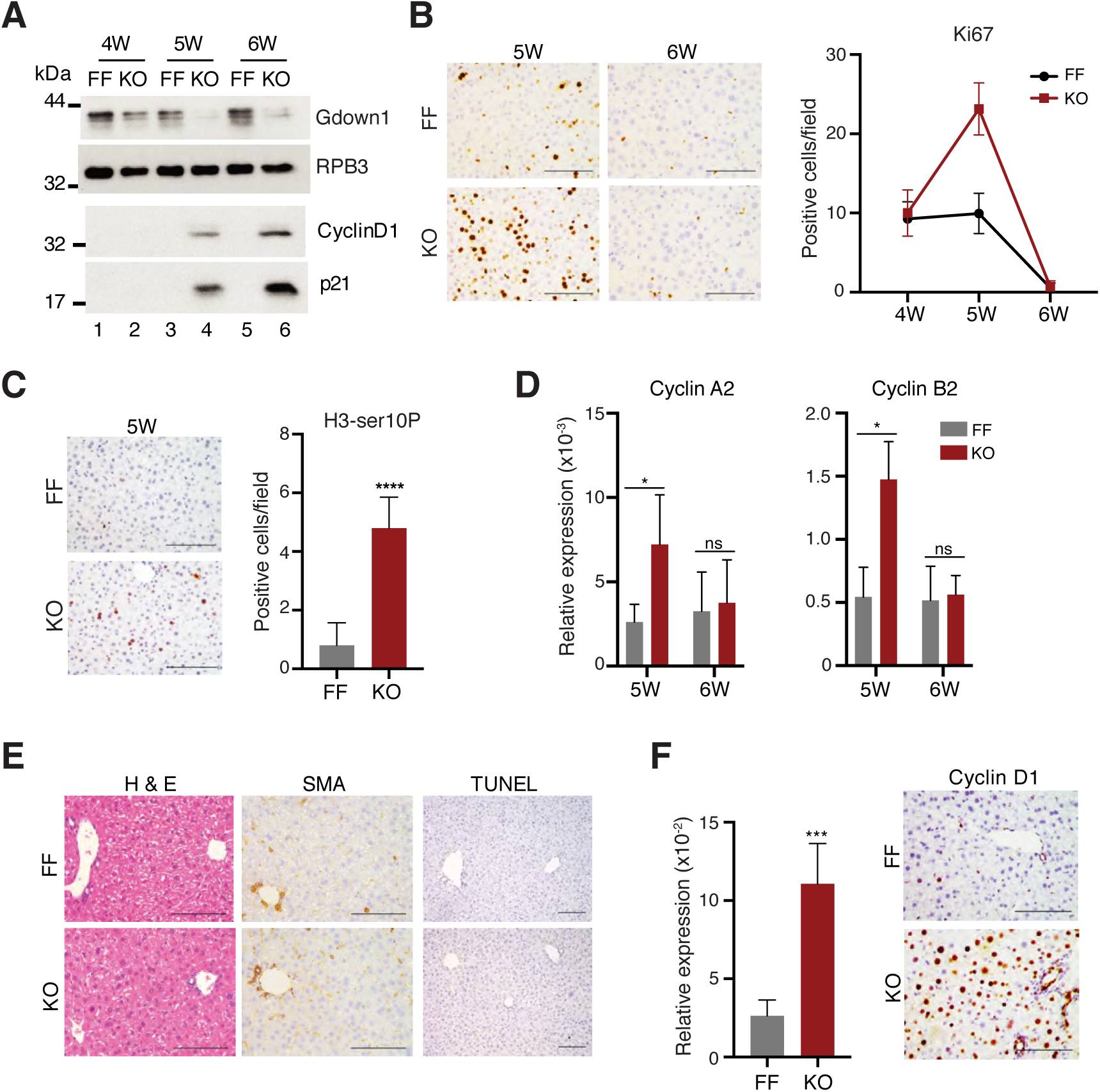
*Gdown1* KO hepatocytes re-enter the cell cycle. (A) Protein expression in control (FF) or Gdown1 liver KO mice (KO) at the indicated ages in weeks (W). Liver whole cell extracts were analyzed by immunoblot. RPB3 serves as a loading control. (B) Left: representative liver histology for Ki67-positive hepatocytes in FF and KO mice at the indicated ages. All scale bars represent 100 μm. Right: quantification of Ki67-positive cells at the indicated ages of weeks (W). (C) Left: representative liver histology for histone H3 serine10-phosphorylated positive hepatocytes in FF and *Gdown1* KO at 5W. All scale bars represent 100 μm. Right: quantification of histone H3 serine10-phosphorylated-positive cells at 5W. \*\*\*\**P* < 0.0001 in unpaired two-tailed t-test. (D) Relative mRNA expression of the indicated genes analyzed by Real-time qPCR. Data are presented with mean and SD (n = 3-4 mice per group at 5W or 6W). \**P* < 0.05 in unpaired two-tailed t-test. (E) Representative liver histology in FF and *Gdown1* KO mice at the ages of 5W. All scale bars represent 100 μm. (F) Left: Relative mRNA expression of cyclin D1 at 6W analyzed by Real-time qPCR. Data are presented with mean and SD (n = 4-5 mice per group). \*\*\**P* < 0.001 in unpaired two-tailed t-test. Right: representative liver histology for cyclin D1-positive hepatocytes in FF and *Gdown1* KO mice at 6W.

Taken together, these data suggest that *Gdown1* KO causes cell cycle re-entry of hepatocytes in the absence of apparent hepatic injury or loss, but that an associated induction of *p21* leads to a subsequent cell cycle arrest.

### *Gdown1* KO causes dysregulated cell cycle progression in the absence of p53

As discussed above, the cell cycle re-entry elicited by *Gdown1* KO in hepatocytes appears to be reversed by *p21* induction, and this appears to be mediated through p53 activation based on the concomitant induction of several known p53 target genes. Notably, *Gdown1* KO hepatocytes were subject to apoptosis at 8W under conditions where the induced p21 would be expected to act as an anti-apoptotic factor. These results suggest that the actual *Gdown1* KO impact on hepatocytes might be partly concealed by the action of the induced p53. To test this possibility, we generated *Gdown1****^f/f;Alb-Cre^*** mice that also carry a *p53* null mutation (designated DKO mice). As expected, *p21* induction was abolished in the DKO liver at 6W (Fig. 3A). Consequently, Ki67-positive DKO hepatocytes, were observed at 6W -- in contrast to what was observed in individual *p53* KO or *Gdown1* KO livers at 6W (Fig. 3B and Fig. 2B). These results indicate that the observed cell cycle re-entry at 5W was subsequently countered by p53-induced p21. Moreover, TUNEL assays failed to detect any apoptotic cells (Fig. 3B), suggesting that this re-entry was not likely to be caused by compensatory proliferation.

**Figure 3.**
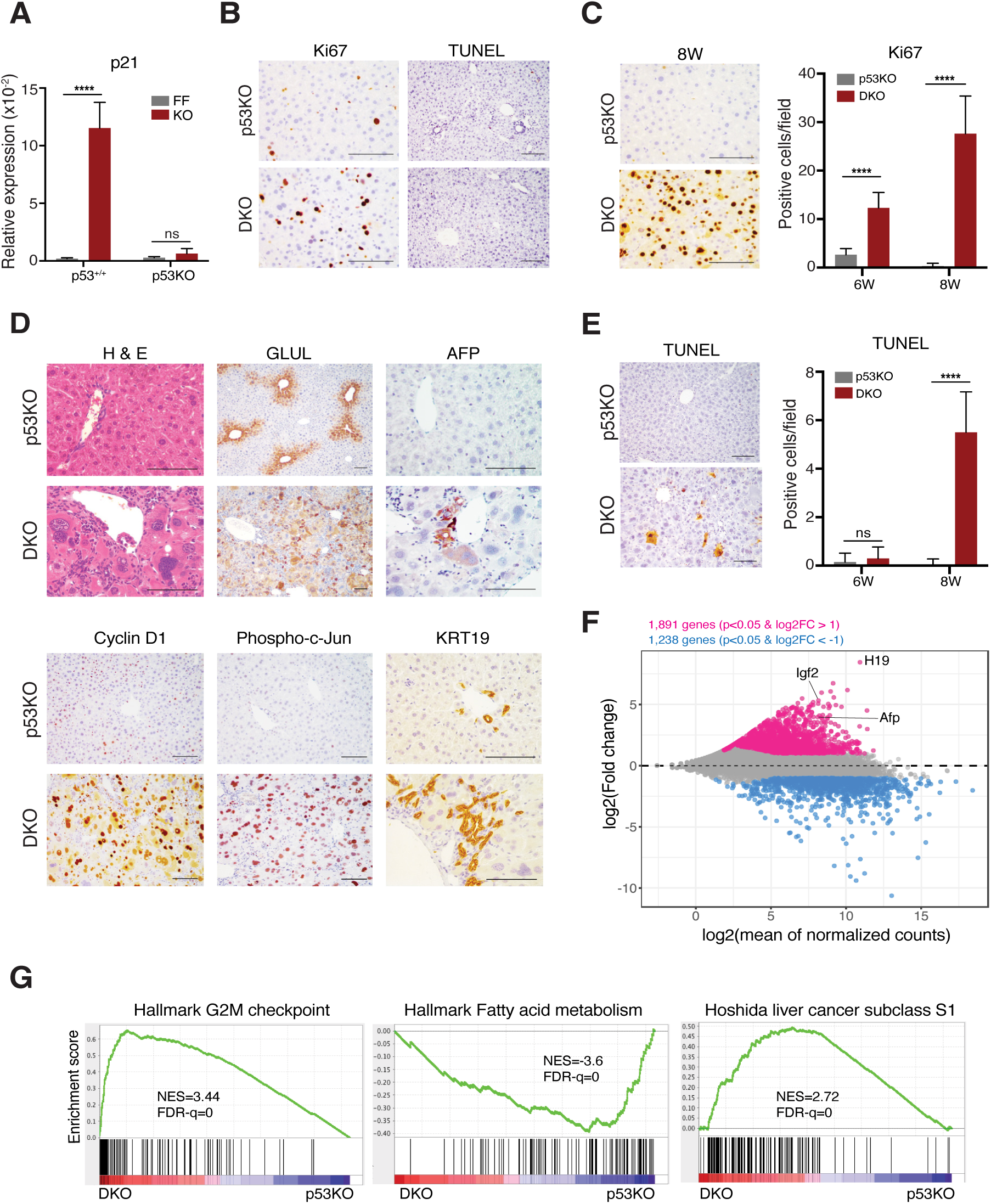
*Gdown1* KO causes dysregulated cell cycle progression in the absence of p53. (A) Relative mRNA expression of p21 in the liver from hepatocyte-specific *Gdown1* KO mice carrying *p53* KO allele (DKO) liver in comparison with *p53* KO liver analyzed by Real-time qPCR. Data are presented with mean and SD (n = 3-5 mice at the ages of 6W per group. \*\*\*\**P* < 0.0001 in unpaired two-tailed t-test. (B) Representative liver histology for Ki67- or TUNEL-positive hepatocytes in FF carrying *p53*KO allele (p53KO) and in DKO mice at 6W. All scale bars represent 100 μm. (C) Left: representative liver histology for Ki67-positive hepatocytes in p53KO and DKO mice at 6W. Right: quantification of Ki67-positive cells at the indicated ages (W). (D) Representative liver histology in p53KO and DKO mice at 8W. (E) Left: representative liver histology for TUNEL-positive hepatocytes in p53KO and DKO mice at 8W. All scale bars represent 100 μm. Right: quantification of TUNEL-positive cells at the indicated ages (W). (F) MA (minus average) plot for differentially expressed genes in DKO liver at 8W. Deep pink or light blue color indicates significantly up- or down-regulated genes, respectively (P < 0.05 and log_2_fold change > 1 or < −1). (G) Gene set enrichment analysis (GSEA) for differentially expressed genes in DKO liver.

At 8W, the number of Ki67-positive DKO hepatocytes was further increased (Fig. 3C). More interestingly, H&E staining showed that the DKO hepatocytes were significantly larger than the normal hepatocytes at this stage, with some being extremely enlarged and containing more than 3 nuclei (Fig. 3D), indicative of incomplete mitoses. Normal liver architecture was obviously distorted, as the pericentral expression pattern of glutamine synthetase (GLUL) was no longer evident in DKO liver (Fig. 3D). Some cells expressed a fetal liver marker, alpha-fetoprotein (AFP), while the expression of cyclin D1 and phosphorylated c-JUN were observed in a majority of the cells (Fig. 3D). The number of apoptotic cells was dramatically increased at this stage (Fig. 3E), which was accompanied by the proliferation of KRT19-positive cells (Fig. 3D). These results show that, in the absence of p53, *Gdown1* KO causes dysregulated cell cycle progression.

To further investigate the DKO phenotype, transcriptome analysis by RNA-seq was performed. The results showed 1891 up- and 1238 down-regulated, genes (Fig. 3F). In particular, the expression of mouse fetal liver markers that include *H19*, *Igf2*, and *Afp* was highly up-regulated (Fig. 3F). GO analysis revealed that the up-regulated genes are enriched in cell cycle, cell adhesion, and immune system processes, while the down-regulated genes are involved in metabolic pathways, particularly in lipid metabolic processes (Supplemental Fig. S3A). The up-regulation of genes, including Aurora kinase genes, that are involved in mitotic nuclear division (Supplemental Fig. S3B) was particularly prominent, which might explain the incomplete mitoses in the DKO phenotype (Fig. 3D). Gene Set Enrichment Analysis (GSEA) further supports the view that the up-regulated genes are enriched in G2M checkpoint genes, E2F target genes, and genes associated with the epithelial-mesenchymal transition (EMT), while the down-regulated genes are enriched in genes associated with fatty acid metabolism (Fig. 3G; Supplemental Fig. S3C). GSEA also revealed that the DKO transcriptome profile is correlated with gene signatures of hepatocellular carcinoma (Acevedo et al., 2008; Borlak et al., 2005; Cairo et al., 2008; Hoshida et al., 2009; Lee et al., 2004; Villanueva et al., 2011) (Fig. 3G; Supplemental Fig. S3D). In particular, the profile is similar to that of human HCC subtype S1 tumors, which exhibit more vascular invasion and satellite lesion properties and whose clinical phenotypes show greater risks of earlier recurrence. Also, the molecular pathways that are characteristic of the tumor relate to activation of the WNT pathway in the absence of β-catenin mutations and up-regulation of TGF-β target genes (Hoshida et al., 2009).

Taken together, these results indicate that the *Gdown1* KO-induced cell cycle progression is subsequently inhibited by an associated *Gdown1* KO induction of p21 through a p53 pathway. In the absence of p53, cell cycle progression is dysregulated in Gdown1-deficient cells, which show a pre-malignant-like transformation. These results suggest that Gdown1 plays a critical role in maintaining normal liver function.

### Gdown1 is associated with elongating Pol II on genes that are actively transcribed in the liver

To investigate how the loss of *Gdown1* causes hepatocytes to re-enter the cell cycle, and for further insights into normal Gdown1 functions, we tried to identify genes that are directly regulated by Pol II(G). To this end, we first examined whether Gdown1 is associated with Pol II in the form of Pol II(G) in mouse liver. Biochemical analyses (Supplemental Fig. S4A) showed that the majority of Gdown1 in whole cell extracts treated with NUN buffer containing 1 M urea, and purified over an anion exchange chromatography (DEAE), was associated with Pol II (monitored by RPB3), while only a very low amount of Pol II-free Gdown1 was isolated by cation exchange chromatography (P11). The result shows that there is little Pol II-free Gdown1 in mouse liver, indicating that Gdown1 plays a direct role in gene transcription through Pol II(G).

To identify genes that are directly targeted by Pol II(G), we performed ChIP-seq analysis with non-crosslinked chromatin prepared from FF (control) or *Gdown1* KO liver at 7W of age. Surprisingly, significant Gdown1 enrichment was observed in gene bodies of highly expressed genes in the liver (Fig. 4A). Among these genes, *Alb*, *Serpina3k*, and *Apob* encode plasma proteins that are constitutively produced in the liver. *Cyp2e1* encodes a member of the cytochrome p450 family that is involved in xenobiotic metabolism, while *Cps1* encodes a mitochondrial enzyme involved in the urea cycle. These genes play a major role in the maintenance of normal liver functions. Notably, the anti-Gdown1 ChIP signals seen in control FF liver were not detected in *Gdown1* KO liver, showing that the observed enrichment signals are Gdown1-specific (Fig. 4A). Also, weaker Gdown1 signals were seen around the promoter-proximal regions compared to the gene bodies (Supplemental Fig. S4B), in contrast to what was previously reported in studies lacking Gdown1 knockout or knockdown cell controls (Cheng et al., 2012). Notably, in knockout cells the Pol II signals on direct Gdown1 target genes were substantially decreased (Fig. 4A, B), whereas Pol II enrichment was increased on genes, such as *Cdo1*, and *Arg1,* that were not directly targeted by Gdown1 (Fig. 4B). The expression of immediate early genes such as *Jun*, *Fos*, and *Btg2* is induced in the priming phase of liver regeneration (Mohn et al., 1990; Thompson et al., 1986). Although these genes are regulated by paused Pol II at their promoter proximal regions (Liu et al., 2015), Gdown1 was not detected on these genes (Fig. 4C). Importantly, Gdown1 also was not observed on genes that are involved in cell cycle progression (Fig. 4C), indicating that *Gdown1* KO-induced cell cycle re-entry is clearly through secondary (indirect) effects.

**Figure 4.**
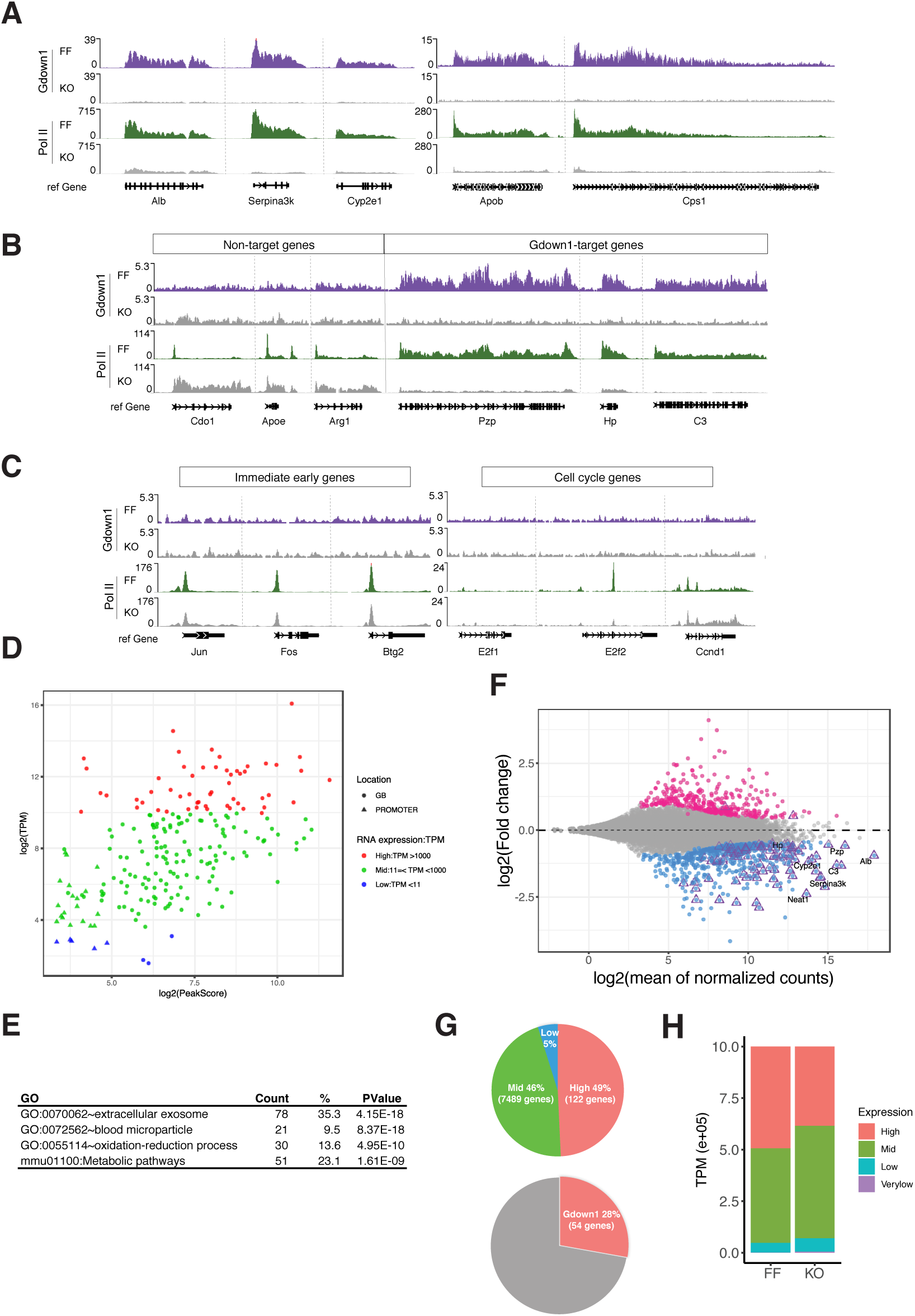
Gdown1 is associated with elongating Pol II on genes that are actively transcribed in the liver. (A, B, C) ChIP-seq profiles generated with Gdown1 and RPB3 (Pol II) antibodies at the indicated genes in FF and *Gdown1* KO liver. (D) Gdown1 target genes that were plotted by transcripts per million (TPM) and ChIP-seq Peak score. (E) Gene ontology analysis for Gdown1 target genes. (F) MA plot for differentially expressed genes in *Gdown1* KO liver at the ages of 7W. Deep pink or light blue color indicates significantly up- or down-regulated genes, respectively (P < 0.05 and log_2_fold change > 0 or < 0). Gdown1 target genes are indicated with triangles in purple color. (G) Top: the percentages of transcripts (TPM) for high-expressed genes (TPM > 1000, shown in red), mid-expressed genes (11 =< TPM < 1000, shown in green), and low-expressed genes (TPM < 11, shown in blue) in FF liver. Bottom: the percentage of transcripts for Gdown1-targeted, highly-expressed genes in FF liver. (H) The percentages of transcripts (TPM) in *Gdown1* KO liver for high-, mid-, low-expressed genes groups that were categorized in FF liver. High: TPM > 1000, Mid: 11 =< TPM < 1000, Low: 1=< TPM < 11, Very low: TPM < 1).

As previously reported (Jishage et al., 2012), the number of genes that are directly targeted by Gdown1 appears to be very low, as peak calling analyses identified only 222 such genes. Plotting peak scores of these genes from ChIP-seq analysis with the corresponding transcripts (transcript per million, TPM) from RNA-seq data showed that 54 of the targeted genes are highly expressed genes in the liver (TPM > 1000) (shown in red circles in Fig. 4D). Gdown1 was also observed in the promoter regions of several genes (Supplemental Fig. S4C) that tend to be categorized as lowly expressed genes (shown in triangles in Fig. 4D). GO analysis revealed that the majority of Gdown1-targeted genes are involved in plasma protein synthesis and metabolic pathways (Fig. 4E). Consistent with the decreased Pol II recruitment upon loss of Gdown1, differential expression analysis showed that mRNA expression of the direct Gdown1 target genes was down-regulated in KO liver (indicated by triangle-enclosed circles in Fig. 4F).

In FF liver, transcripts of 122 genes (transcript per million (TPM) > 1000) account for approximately 50% of the entire liver mRNA, with about 44% of the corresponding genes being occupied by Pol II(G) (Fig. 4G). Although the number of genes that were identified as Pol II(G) direct target genes seems to be quite low (less than 0.4% of the total number of expressed genes), approximately 30% of total mRNA synthesis is derived from genes directly regulated by Pol II(G) transcription (Fig. 4G). Also, TPM count analysis showed that the total transcripts of highly expressed genes (TPM > 1000) were decreased in the KO, while the remaining gene transcripts were significantly increased (Fig. 4H).

In summary, the majority of Gdown1 is detected in association with elongating Pol II on direct target genes that are important for maintaining normal liver functions, suggesting that Gdown1 is directly involved in regulation of the expression of select genes.

### Expression of direct Gdown1 target genes is inversely correlated with expression of cell cycle-related genes

Although *Gdown1* KO leads initially to cell cycle re-entry, the nature of the down-regulated genes in the KO hepatocytes indicates that Gdown1 is not directly involved in transcription of the up-regulated cell cycle-related genes. Rather, the down-regulated expression of Gdown1 direct target genes (identified by the ChIP-seq) seems to be the primary effect of the *Gdown1* KO relating to cell cycle re-entry. However, since the *Gdown1* KO resulting from *Alb-Cre* expression is complete by 5 weeks, this down-regulation of Gdown1 direct target gene expression in the KO could be due to secondary effects. Therefore, to investigate the immediate effects of the *Gdown1* KO, we generated tamoxifen-inducible *Gdown1****^f/f^*** (KO) mice carrying both the *Alb-CreERT2* transgene and a *p53* null mutation (designated DKO***^Alb-CreERT2^*** mice). Around 6 hours after tamoxifen injection, the level of Gdown1 protein relative to the level of the RPB3 subunit of Pol II was significantly decreased, and by 14 hours declined to nearly undetectable levels (Fig. 5A).

**Figure 5.**
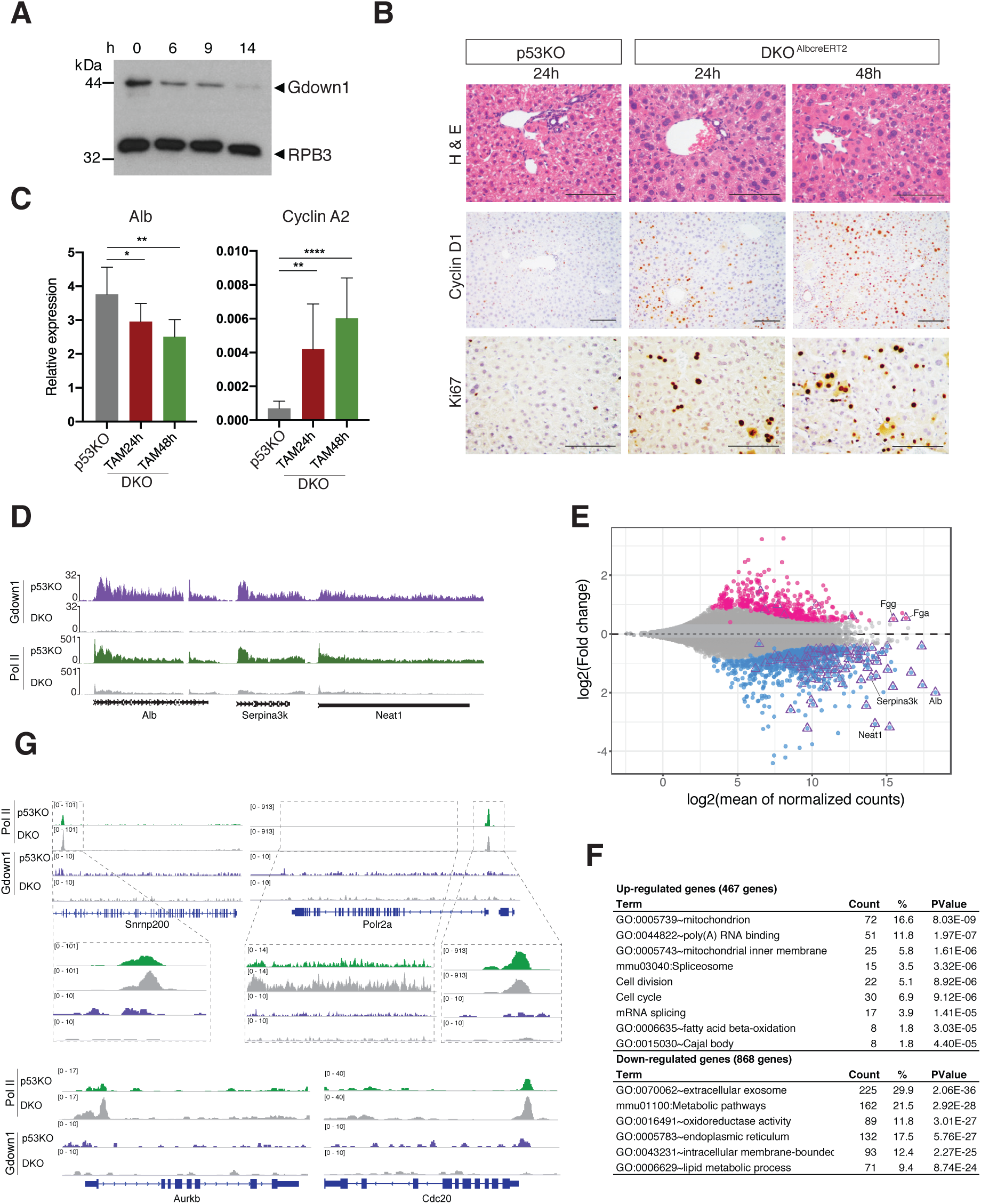
Expression of Gdown1 target genes is inversely correlated with expression of cell cycle-related genes. (A) Gdown1 expression in liver after tamoxifen injection in FF mice carrying both the *p53* KO allele and *Alb-creERT2* (DKO***^Alb-creERT2^*** mice) at the indicated time (h: hour). Liver nuclear pellets were analyzed by immunoblot. RPB3 serves as a loading control. (B) Representative liver histology in *p53* KO and DKO***^Alb-creERT2^*** mice at the indicated times. All scale bars represent 100 μm. (C) Relative mRNA expression of albumin (*Alb*) and *cyclin A2* genes analyzed by Real-time qPCR. Control, *p53* KO mice treated with tamoxifen for 24 hours (n = 9). TAM24h and TAM48h, DKO***^Alb-^*^creERT2^** mice treated with tamoxifen for 24 or 48 hours (n = 7 or 5), respectively. Data are presented with mean and SD. \**P* < 0.05, \*\**P* < 0.01, \*\*\*\**P* < 0.0001 in unpaired two-tailed t-test. (D) ChIP-seq profiles generated with Gdown1 and RPB3 (Pol II) antibodies at the indicated genes in p53KO and DKO***^Alb-creERT2^*** liver. (E) MA plot for differentially expressed genes in DKO***^Alb-creERT2^*** liver treated with tamoxifen for 24 hours. Deep pink or light blue color indicates significantly up- or down-regulated genes, respectively (P < 0.05 and log_2_fold change > 0 or < 0). Direct Gdown1 target genes are indicated with triangle-enclosed circle. (F) Gene ontology analysis for differentially expressed genes in DKO***^Alb-creERT2^*** liver. (G) ChIP-seq profiles generated with Gdown1 and RPB3 (Pol II) antibodies at the indicated genes in p53KO and DKO (DKO***^Alb-creERT2^***) liver.

Histological analyses showed that DKO***^Alb-CreERT2^*** hepatocyte nuclei were enlarged relative to *p53* KO hepatocyte nuclei (H&E staining in Fig. 5B). Cyclin D1 expression was detected in some DKO***^Alb-CreERT2^*** hepatocytes within 24 hours after tamoxifen injection and was observed in most hepatocytes at 48 hours (Fig. 5B). Similarly, significant numbers of DKO***^Alb-creERT2^*** hepatocytes were Ki67-positive within 24 hours (Fig. 5B). These results show that DKO***^Alb-creERT2^*** hepatocytes re-enter the cell cycle quite rapidly.

Consistent with the results observed in the *Gdown1****^f/f;Alb-Cre^*** (KO) hepatocytes (Fig. 4F), albumin mRNA expression in DKO***^Alb-creERT2^*** hepatocytes gradually decreased within 48 hours after tamoxifen injection, while cyclin A2 expression dramatically increased (Fig. 5C). To further analyze the inversely correlated expression, we performed ChIP-seq combined with RNA-seq using liver from DKO***^Alb-creERT2^*** mice treated with tamoxifen for 24 hours. ChIP-seq profiles showed that the *Gdown1* KO led to decreased Pol II occupancies on the direct target genes (Fig. 5D). Consistent with the results observed in KO hepatocytes (Fig. 4F), RNA-seq analysis in DKO***^Alb-creERT2^*** hepatocytes showed that expression of a majority of the direct Gdown1 target genes (purple triangle-enclosed circles in Fig. 5E) were down-regulated upon loss of Gdown1. Noted exceptions were eight genes, including *Fgg* and *Fga* (indicated in Fig. 5E), whose expression was moderately up-regulated (less than 1.5-fold). Consistent with these RNA-seq results, ChIP-seq profiles showed that the Pol II signal on the *Fgg* gene body was unaffected even though Gdown1 was no longer detected (Supplemental Fig. S5A). However, Pol II recruitment on this gene was decreased in KO***^Alb-cre^*** hepatocytes at 7W (Supplemental Fig. S5B), which suggests that the kinetics of an impact of Gdown1 loss on Pol II recruitment may vary depending on the specific gene. GO analysis for differentially expressed genes in DKO***^Alb-creERT2^*** hepatocytes showed results similar to those observed (Supplemental Fig. S3A) for the down-regulated genes in DKO***^Alb-Cre^*** hepatocytes, with the up-regulated genes being particularly enriched not only in cell cycle-related genes but also in genes related to mitochondrial and RNA metabolism (Fig. 5F; Supplemental Fig. S5C).

In relation to the up-regulated genes in DKO***^Alb-CreERT2^*** hepatocytes, the *Snrnp200* gene that encodes the small nuclear ribonucleoprotein U5 subunit 200 is one of the up-regulated spliceosome genes indicated by GO analysis (Supplemental Fig. S5C). The ChIP-seq profile clearly shows increased Pol II recruitment to the promoter region in DKO***^Alb-creERT2^*** hepatocytes relative to control (*p53* KO) hepatocytes (Fig. 5G). Similar Pol II ChIP-seq profiles, with increased Pol II recruitment to promoter regions, were also observed for the up-regulated cell cycle-related *Aurkb* and *Cdc20* genes (Fig. 5G). The up-regulated *Polr2a,* encoding the RPB1 subunit of Pol II, showed a reduction in the level of a strong paused Pol II at the promoter and a reciprocal increase in elongating Pol II in DKO***^Alb-creERT2^*** hepatocytes relative to *p53* KO hepatocytes (Fig. 5G). As predicted, Gdown1 was not detected on these up-regulated genes in *p53* KO hepatocytes, indicating that their up-regulation upon Gdown1 loss is indeed through indirect effects of Gdown1.

Since Pol II recruitment to direct Gdown1 target genes rapidly decreases in the DKO***^Alb-CreERT2^*** liver, the normal role of Gdown1 on Pol II in the bodies of these genes remains unclear. However, there are several genes whose transcripts were increased in DKO***^Alb-CreERT2^*** liver by a seemingly failed termination of transcription of an adjacent Gdown1 target gene, as exemplified by the following. Expression of *Sftpa1*, which encodes the surfactant associated protein A1, is normally restricted to adult lung but increased in DKO***^Alb-CreERT2^*** liver (Supplemental Fig. S5D). Interestingly, on *Mat1a*, a down-regulated direct Gdown1 target gene that is located just up-stream of *Sftpa1* (Supplemental Fig. S5E), Pol II was detected continuously from the termination site of *Mat1a* to the gene body of *Sftpa1* in DKO***^Alb-CreERT2^*** liver, implying a potential role for Gdown1 in transcription termination of *Sftpa1* and read-through of the downstream gene in the absence of Gdown1.

Altogether, the results indicate that *Gdown1* KO can rapidly (within 24 hours) stimulate hepatocytes to re-enter into the cell cycle in the absence of p53. Notably, the down-regulated expression of direct Gdown1 target genes upon Gdown1 loss appears to be inversely correlated with the expression of genes that are involved in cell cycle control and mitochondrial and RNA metabolism.

### Down regulation of highly expressed genes in the liver proceeds the cell cycle re-entry

Although the tamoxifen-inducible *Gdown1* KO results in DKO***^Alb-CreERT2^*** liver clearly show an inverse correlation between the expression of direct Gdown1 target genes and cell cycle-related genes, it remains unclear how the DKO hepatocytes can re-enter into the cell cycle. To investigate the basis for the re-entry, we analyzed DKO***^Alb-CreERT2^*** liver at an earlier time point (16 hours) after tamoxifen administration. Notably, two distinct groups of DKO***^Alb-CreERT2^*** livers were distinguished with respect to the level of cyclin D1 RNA expression (indicated as ‘High” and “Low” in Fig. 6A), while *Alb* RNA expression was significantly down-regulated in most all cases (Fig. 6A). Histological analyses further indicated the absence of cyclin D1- and Ki67-positive cells in DKO***^Alb-CreERT2^*** livers with the low level of cyclin D1 RNA expression (indicated as “Low” in Fig. 6B). For DKO***^Alb-CreERT2^*** livers with the high level of cyclin D1 RNA expression, there were sub-groups that were either cyclin D1 positive and Ki67 negative or both cyclin D1 positive and Ki67 positive (identified as “High” in Fig. 6B), indicating normal cell cycle progression in which cyclin D1 expression proceeds prior to Ki67 expression. ChIP-qPCR analyses further revealed that the amounts of Pol II on the bodies of at least two genes, the *Alb* and *C3* genes that are direct Gdown1 target genes, were significantly decreased in the group with the low level of cyclin D1 RNA expression (Fig. 6C). This result shows that the down-regulation of these direct Gdown1 target genes occurs prior to cyclin D1 expression. However, not all the target genes were down-regulated simultaneously. For *Serpina3k,* one of the direct Gdown1 target genes, the decrease of Pol II in the gene body was not observed in the group with the low level of cyclin D1 but was observed in the group with the high level of cyclin D1 (Fig. 6C), suggesting that the time it takes to see an impact of Gdown1 loss on Pol II recruitment is gene-dependent.

**Figure 6.**
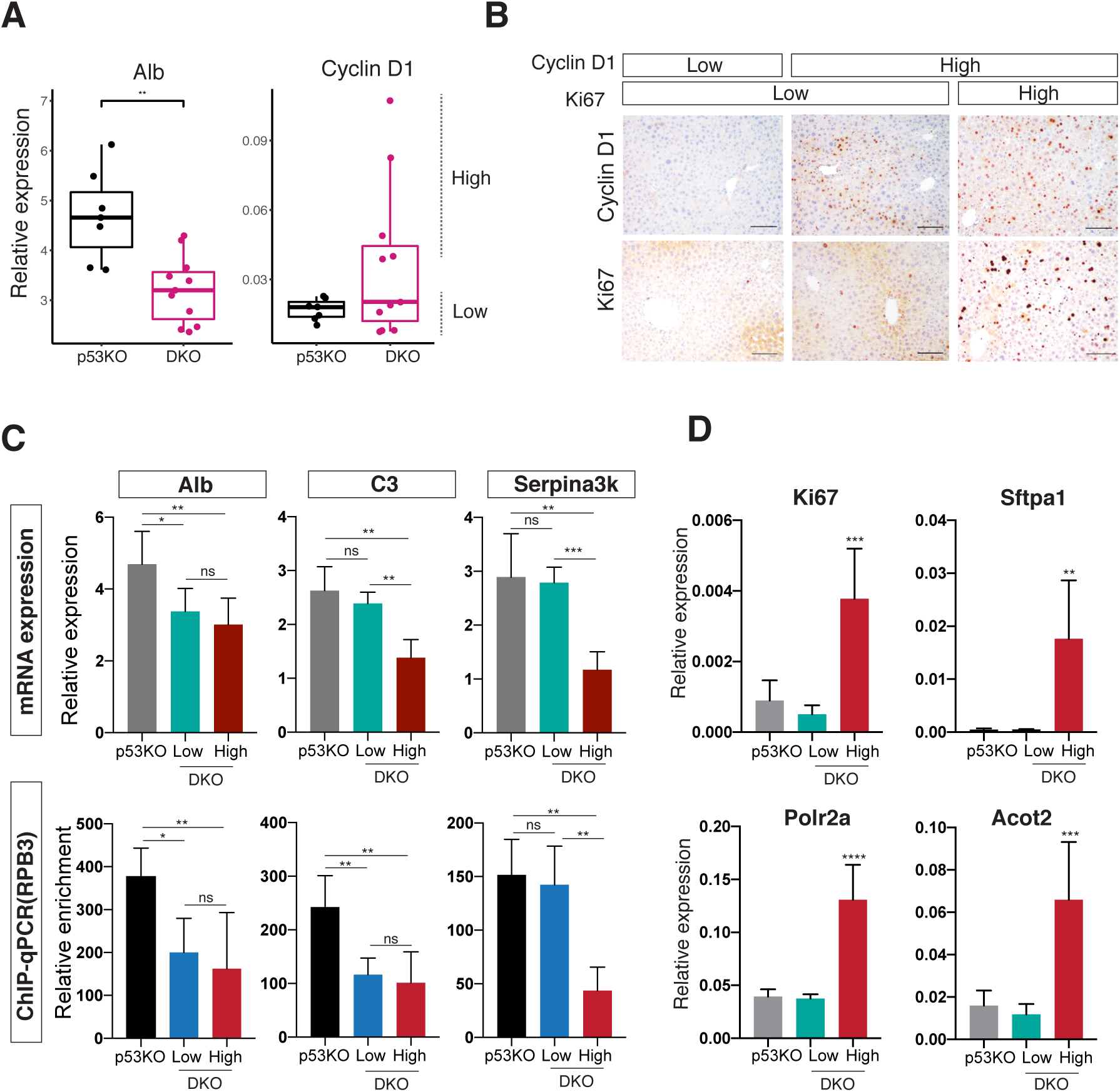
Down regulation of highly expressed genes in the liver induces cell cycle re-entry. (A) Relative mRNA expression of albumin (*Alb*) and *cyclin D1* genes analyzed by Real-time qPCR. Control, *p53* KO mice treated with tamoxifen for 16 hours (n = 7). DKO, DKO***^Alb-creERT2^*** mice treated with tamoxifen for 16 (n = 11), respectively. \*\**P* < 0.01, in unpaired two-tailed t-test. Two distinct groups of DKO***^Alb-creERT2^*** livers, in terms of the level of cyclin D1 RNA expression, are indicated as ‘High” (n=5) and “Low” (n=6). (B) Representative liver histology in DKO***^Alb-creERT2^*** mice expressing high or low levels of cyclin D1 and Ki67. All scale bars represent 100 μm. (C) Top: relative mRNA expression of the indicated genes analyzed by Real-time qPCR. Bottom: relative enrichment of Pol II on the gene bodies for the indicated genes analyzed by Real-time qPCR. Control, *p53* KO mice treated with tamoxifen for 16 hours (n = 7 mice per group). Low or High, DKO***^Alb-creERT2^*** mice treated with tamoxifen for 16 hours and expressing low or high levels of cyclin D1 (n = 6 or 5 mice per group). Data are presented with mean and SD. \**P* < 0.05, \*\**P* < 0.01, \*\*\*\**P* < 0.0001 in unpaired two-tailed t-test. (D) Relative mRNA expression of the indicated genes analyzed by Real-time qPCR. Control, *p53* KO mice treated with tamoxifen for 16 hours (n = 7 mice per group). Low or High, DKO***^Alb-creERT2^*** mice treated with tamoxifen for 16 hours and expressing low or high s of cyclin D1 (n = 6 or 5 mice per group). Data are presented with mean and SD. \*\**P* < 0.01, \*\*\*\**P* < 0.0001 in unpaired two-tailed t-test.

Genes that might be induced by *Gdown1* KO prior to activation of cyclin D1 expression could be involved in the direct cause of the cell cycle re-entry. Therefore, RNA expression levels were assessed for several genes that were up-regulated in DKO***^Alb-CreERT2^*** liver at 24 hours. However, no induction of these genes was observed in the liver group with low levels of cyclin D1 (Fig. 6D). We also analyzed expression of immediate early genes whose expression is induced at the early stage of liver regeneration, although no induction of four such genes (*Myc*, *Jun*, *Fos*, and *Btg2*) was observed in the group with the low levels of cyclin D1 RNA expression (Supplemental Fig. S6A).

As previously reported, overexpression of cyclin D1 is sufficient to induce hepatocyte proliferation in vivo (Mullany et al., 2008; Nelsen et al., 2001). As the *Gdown1* KO-induced cell cycle re-entry could simply be stimulated by the induction of cyclin D1 expression, we investigated the mechanism of the induction. Cyclin D1 expression is regulated by both transcriptional and post-transcriptional mechanisms (Witzel et al., 2010). ChIP-qPCR analysis showed that Pol II recruitment was increased at the promoter region of *cyclin D1* in DKO***^Alb-CreERT2^*** liver with the high cyclin D1 RNA expression (Supplemental Fig. S6B), indicating that the *cyclin D1* induction involves transcriptional activation. It is known that cyclin D1 expression is mitogen-activated, and its promoter is regulated by multiple transcription factors (Klein and Assoian, 2008) that include STAT3, c-JUN, NF-κB, and β-catenin. Although it appears that there is no obvious liver damage that might activate a mitogenic signaling pathway in DKO***^Alb-CreERT2^*** liver, the nuclear localization of these transcription factors was analyzed by immunoblotting of nuclear pellets. Whereas acute injury by carbon tetrachloride induced the expression of β-catenin, phosphorylated STAT3, and NF-κB(p65) in the nucleus at 12 h after administration (Supplemental Fig. S6C, lane 9 versus lanes 10 and 11), β-catenin induction was not observed in the DKO***^Alb-CreERT2^*** liver (lanes 4-8 versus lane 9 or lanes 10 and 11). Nuclear-localized phosphorylated STAT3, as well as NF-κB, was detected in several DKO***^Alb-CreERT2^*** livers (Supplemental Fig. S6C, lanes 5, 6, and 8). However, expression levels were not proportionally correlated with cyclin D1 mRNA expression levels, such that the immediate molecular basis for cyclin D1 induction in DKO***^Alb-CreERT2^*** liver remains to be determined.

### Expression of highly expressed liver-specific genes are inversely correlated with expression of cell cycle-related genes in hepatocellular carcinoma

Although the mechanism of *cyclin D1* induction remains unknown, the down-regulation of Gdown1-associated genes, especially those highly expressed in the liver, appears to be correlated with cell cycle re-entry. Supporting this idea, a differential expression analysis of a TCGA hepatocellular carcinoma (HCC) cohort with high-impact p53 mutations from public patient data sets revealed an inverse correlation between the expression of cell cycle-involved genes and highly expressed genes in the liver (Fig. 7A). Compared to the normal liver tissue (Cluster1 in Fig. 7A), the inverse correlation is quite evident for Cluster 2, while Clusters 3 and 4 show partial correlation (Fig. 7A; Supplemental Fig. S7A). Homeobox genes and tumor antigens were highly expressed in the HCC cohort. However, an inverse correlation of these genes with highly expressed genes in the liver is less clear than the inverse correlation of the highly expressed genes with cell cycle-involved genes (Supplemental Fig. S7B, C). CENPF is a kinetochore protein (Liao et al., 1995), whose overexpression is frequently observed in various cancers including HCC (Dai et al., 2013). The inverse correlation between albumin and CENPF is more convincing compared to a homeobox protein, HOXA9 (Fig. 7B).

**Figure 7.**
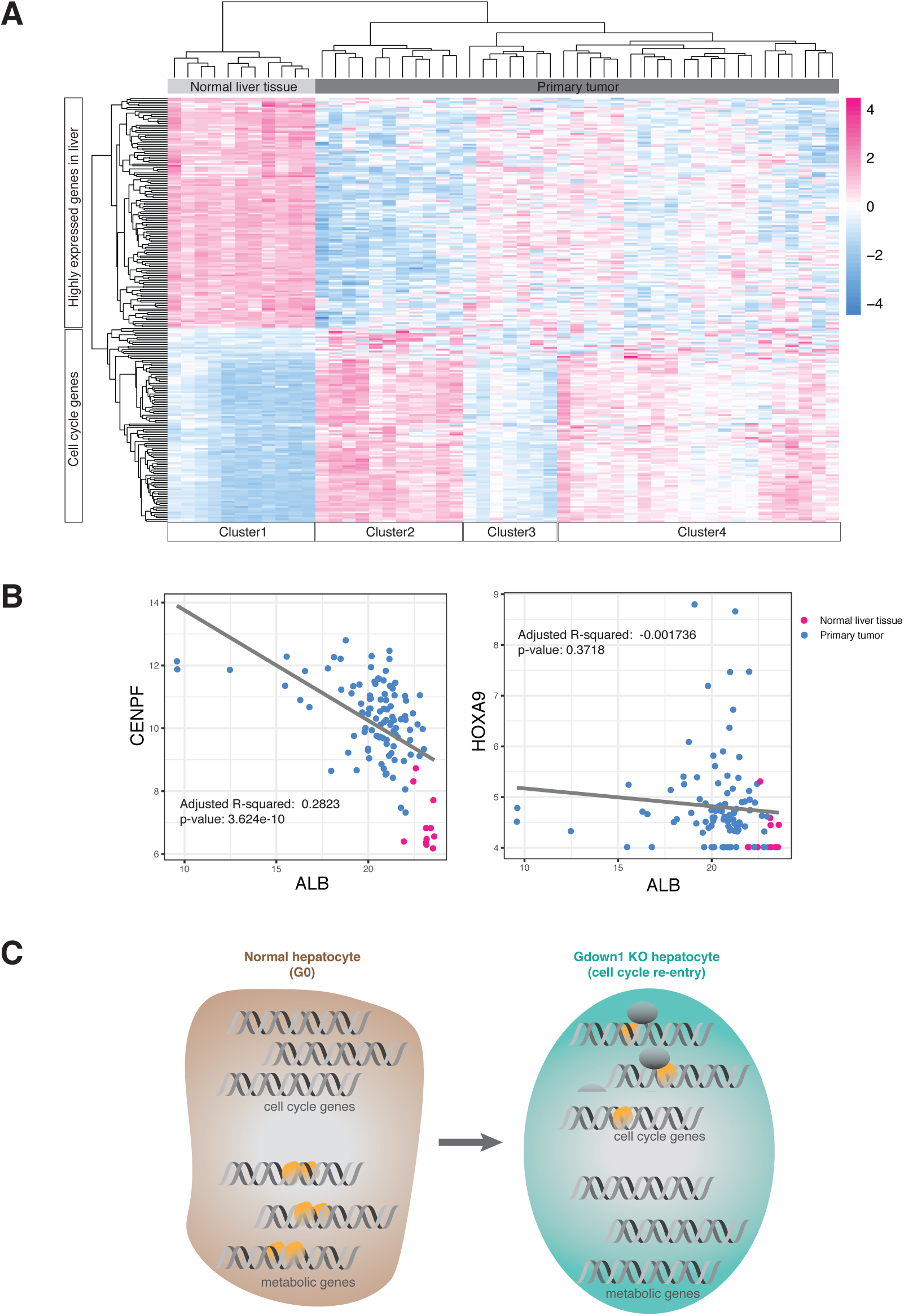
Expression of metabolic genes is inversely correlated with expression of cell cycle-related genes in HCC. (A) Comparison of mRNA expression of cell cycle-related genes and highly expressed liver-specific genes in normal liver tissue or in HCC cohorts with high impact p53 mutations. (B) Scatter plot and regression line data for expression of *ALB* and *CENPF* or *HOXA9* in normal liver tissue (shown in pink) and in HCC cohorts with p53 mutations (shown in blue). (C) Schematic of Pol II recruitment to gene regions in the presence or absence of Gdown1.

Overall, these latter results suggest that the down-regulation of highly expressed genes in the liver may contribute to hepatocyte proliferation leading to tumor development.

## Discussion

In this study, we report a direct role of the Pol II-associated factor Gdown1 in the regulation of gene transcription to maintain normal liver functions. Gdown1 is bound to Pol II as Pol II(G) in the liver and localized to the gene bodies of highly transcribed genes. The loss of Gdown1, through a hepatocyte-specific *Gdown1* knockout (KO), leads to the down-regulation of these genes, which in turn activates expression of cell cycle regulatory genes and leads to hepatocyte re-entry into the cell cycle. However, this cell cycle re-entry is ultimately countered by a *Gdown1* KO-mediated, p53-dependent induction of p21-- as revealed by a facilitated cell cycle entry and resultant cell cycle dysregulation with a double knockout of *Gdown1* and *p53*. Most importantly, these results establish an important and newly recognized physiological function for an RNA polymerase II regulatory factor (Gdown1) in the maintenance of normal liver cell transcription through constraints on cell cycle re-entry of quiescent hepatocytes.

Consistent with results of our previous study of Drosophila Gdown1 (Jishage et al., 2018), our finding that *Gdown1* KO mice are embryonic lethal indicates an essential role for Gdown1 in mouse early embryonic development. However, Gdown1 hepatocyte KO mice appear to be normal and healthy, suggesting that the loss of Gdown1 is not lethal for quiescent cells such as hepatocytes. Although the observed phenotype in the *Gdown*1 KO (KO***^Alb-Cre^***) liver is relatively less severe compared to the *p53;Gdown1* double knockout (DKO***^Alb-Cre^***) liver, the major impact on the liver is the continuous expression of cyclin D1 followed by activation of p53 signaling pathways. Although the cell cycle progression of *Gdown1* KO hepatocytes observed at 5 weeks is subsequently countered by p53-induced p21, a significant number of cells re-enter the cell cycle or succumb to apoptosis by 8 weeks. While p21 arrests the cell cycle by inhibiting cyclin-dependent kinases (CDKs), it also has oncogenic activities (Abbas and Dutta, 2009). In this regard, the direct binding of p21 to cyclin D1 is one way to maintain the nuclear localization of cyclin D1, which promotes the assembly of cyclin D1/CDK4 or cyclin D1/CDK6 complexes without inhibiting the kinase activities (Alt et al., 2002; LaBaer et al., 1997). The observed cell cycle re-entry at 8 weeks might be explained by the p21 oncogenic activity; and the expression of pro-apoptotic genes activated by p53 could cause cell death. Although the exact mechanism that triggers activation of an apoptotic pathway remains unclear, the continuous expression of cyclin D1 may contribute to the driving of hepatocytes to cell death. Supporting this possibility, neuronal programmed cell death is accompanied by cyclin D1 induction, and overexpression of cyclin D1 causes apoptosis not only in neural cells but also in non-neural cells (Freeman et al., 1994; Kranenburg et al., 1996). Also, apart from a role in cell cycle progression, cyclin D1 has several additional functions, including the regulation of transcription factors (Musgrove et al., 2011; Pestell, 2013), that might contribute to the observed apoptosis.

p53 is a tumor suppressor that is most frequently mutated in human cancers (Kastenhuber and Lowe, 2017). The more severe phenotype of the double *Gdown1;p53* KO (DKO***^Alb-cre^***) liver relative to the *Gdown1* KO liver indicates a p53 protective role in the *Gdown1* KO context. Although cell cycle regulators, such as cyclin A2 or cyclin B2, that are required for proper progression are all expressed in DKO***^Alb-cre^*** liver, the cell cycle is clearly dysregulated and generates incomplete mitoses. The mechanism underlying this dysregulation is unknown, although the up-regulated expression of mitotic genes might be involved. Thus, Aurora-A kinase overexpression, which is often observed in various cancers, leads to disruption of the cell cycle checkpoint as well as to aneuploidy (Marumoto et al., 2005). In this regard, Aurora-B, overexpression of which causes effects similar to those of Aurora-A (Willems et al., 2018), was also up-regulated in DKO***^Alb-cre^*** liver. Since Aurora kinases and p53 are mutually regulated (Sasai et al., 2016), the Aurora-B overexpression resulting from p53 loss could cause the cytokinesis failure observed in DKO***^Alb-cre^*** hepatocytes.

Apart from the histological evidence for dysplasia in DKO***^Alb-cre^*** liver, the associated RNA expression profile, including the up-regulation of cyclin D1, is similar to that of several types of hepatocellular carcinoma (HCC) -- suggesting the potential for pre-malignant transformation. *Cyclin D1* amplification and overexpression are frequently observed in a wide variety of cancers (Musgrove et al., 2011). Also, transgenic mice that overexpress cyclin D1 in the liver develop hepatomegaly at 6 months, which leads to the formation of adenomas or carcinomas (Deane et al., 2001). Considering the *Gdown1* KO impact on gene expression, including the cyclin D1 overexpression that was further amplified by p53 ablation, the DKO***^Alb-cre^*** liver might be expected to develop tumors. Although we did not observe malignant transformation in DKO***^Alb-cre^*** liver at 6 months of age, further analysis was prevented by a *p53* KO-caused lymphoma that leads to death around 8 months. Therefore, an alternative approach is necessary to examine this possibility.

In relation to direct transcription functions of Gdown1, in vitro studies have shown (i) an ability of Gdown1 to inhibit transcription by preventing initiation factor TFIIF binding to Pol II (Jishage et al., 2012) and (ii) an ability of Mediator to reverse the transcriptional repressive capacity of Gdown1, rendering Pol II(G) particularly dependent on Mediator to initiate transcription (Hu et al., 2006). Thus, Pol II(G) may not be recruited to (or be active on) gene promoters on which Mediator/activator is not present, such that inappropriate Pol II recruitment to (or activity on) such promoters is restricted by Gdown1. In the liver, the majority of Gdown1 is found in association with Pol II as Pol II(G). Consistent with the mutually exclusive interaction of Gdown1 and TFIIF with Pol II, most of the Gdown1 is not observed either on promoters or at promoter proximal regions. Unexpectedly, however, Gdown1 was detected on the gene bodies of actively transcribed genes in the liver. This raises an important question as to how Gdown1 associates with Pol II during the various phases of the transcription cycle -- from Pol II recruitment to elongation. Our structural studies and in vitro transcription assays with purified factors (Hu et al., 2006; Jishage et al., 2012; Jishage et al., 2018) suggested a model in which Mediator interaction with promoter-associated Pol II(G) de-stabilizes Gdown1 association with Pol II and thus allows TFIIF binding to Pol II. Therefore, it is reasonable to speculate that Gdown1 dissociation from Pol II(G) must occur for transcriptional activation (including initiation), which is generally Mediator-dependent in cells. However, our current results suggest that, at least for the highly active genes in hepatocytes, the Gdown1-Pol II interaction is maintained through the entire transcriptional cycle. One possibility is that the Gdown1-Pol II interaction is dynamic and reformed after Pol II initiation and promoter clearance. Consistent with this idea, biochemical assays on a model promoter have indicated an ability of Gdown1 to re-associate with Pol II after promoter clearance (DeLaney and Luse, 2016). Future biochemical assays with purified factors on relevant genes, occupied by Pol II(G) in cells, should provide further information on this point.

In relation to the Gdown1 post-initiation functions (including termination) indicated here, the specific role of Gdown1 in association with elongating Pol II in gene bodies remains unclear. However, one previous study showed that Pol II(G) does not inhibit Pol II elongation in a purified in vitro assay system (Hu et al., 2006), while another study showed that Gdown1 can actually facilitate Pol II elongation in a less defined in vitro assay (Cheng et al., 2012). Relevant to possible Pol II-associated Gdown1 interactions during elongation, a recent cryo-EM study described an activated Pol II elongation complex that includes DSIF, the PAF1 complex (PAF1C), and SPT6 (Vos et al., 2018); and superimposing the Pol II(G) cryo-EM structure (Jishage et al., 2018) on the elongation complex structure reveals no overlap with these elongation factors (except for LEO1 in PAF1C) in the static structure. However, it remains unknown whether Pol II-bound Gdown1 interacts directly with PAF1C (or other factors) in the elongation complex and how PAF1C regulates elongating Pol II in normal mouse liver. A further understanding of the function and mechanism of action mechanism of Gdown1 through interactions with elongating Pol II will necessitate further biochemical studies with more physiological (chromatin) templates where elongation factor functions are especially evident (Kim et al., 2010).

The loss of Gdown1 in the liver leads to substantial decreases in Pol II association with highly transcribed genes, resulting in the down-regulation of RNA expression. Notably, RNA transcripts of these Gdown1-targeted genes account for approximately 30% of total liver transcripts. Therefore, the impact of the reduction of Pol II recruitment to these genes would not be insignificant for global gene expression. Notably, the Pol II recruitment to the albumin gene is immediately affected by *Gdown1* KO. The albumin gene is the most highly expressed gene in the liver, and its transcriptional activation depends on an enhancer region that lies around 10 kb upstream from the transcription start site (Pinkert et al., 1987) and closely resembles super-enhancers. Super-enhancers are extremely sensitive to perturbation of associated components (Hnisz et al., 2017), such that loss of Gdown1 might contribute to the prompt reduction of Pol II recruitment to the albumin gene.

There is an unambiguous inverse correlation between the expression of Gdown1 target genes and cell cycle-related genes. The down-regulated expression of highly activated genes in the liver leads to the immediate up-regulation of genes encoding proteins involved in poly(A) binding, spliceosome function, and mitochondrion function, which are essential cellular components. Also, *cyclin D1* transcription is activated in an unknown and apparently non-canonical manner. In normal liver, hepatocytes are committed to expressing highly selected genes that are involved in synthesis of plasma proteins or in metabolism. Thus, a disproportionate amount of Pol II is engaged in transcription of these genes. Although the exact mechanism leading to activation of cell cycle-related genes remains unclear, a reduction of Pol II recruitment to the liver-specific genes could make Pol II available to other genes. Also, in the absence of Gdown1, which restricts Mediator/activator–independent transcription, Pol II recruitment could occur in a less competitive environment. Consequently, the expression of heavily Mediator/activator-dependent genes would be decreased, while genes whose transcription is maintained at the basal level might be activated. This Pol II re-allocation to other genes might underlie hepatocyte re-entry into the cell cycle (Fig. 7C).

The recent genomic characterization of HCCs has shown frequent mutations in metabolic genes that include *ALB*, *APOB*, and *CPS1* (Cancer Genome Atlas Research Network. Electronic address and Cancer Genome Atlas Research, 2017; Fujimoto et al., 2016; Schulze et al., 2015); and genomic alterations in *ALB* may down-regulate *ALB* expression (Fernandez-Banet et al., 2014). Previous studies also suggest that metabolic reprogramming plays a key role in the progression of hepatocytes to malignant HCC (Cancer Genome Atlas Research Network. Electronic address and Cancer Genome Atlas Research, 2017). However, the molecular mechanisms that contribute to the tumorigenesis have remained unknown. Our study provides important insights into a mechanism whereby down-regulation of highly expressed genes could contribute to induction of hepatocyte re-entry into the cell cycle, and thus has important implications both for hepatocarcinogenesis and for liver regeneration.

## Materials and methods

### Animals

*Gdown1^flox/flox^* mice were generated using a Gdown1 targeting vector (Supplemental Fig. S1A) that was obtained from the KOMP (the trans-NIH Knock-Out Mouse Project) Repository. *Gdown1^flox/flox^* mice were maintained on a C57BL/6 background. To conditionally delete *Gdown1* in the liver, *Gdown1^flox/flox^* mice were crossed with transgenic mice expressing *Cre* under the control of the albumin promoter (*Alb-Cre* mice) (Stock Number 003574, The Jackson Laboratory). To generate *p53* and *Gdown1* double knockout (DKO*^Alb-cre^*) mice, *p53^-/-^* mice (Stock Number 002101, The Jackson Laboratory) were crossed with *Gdown1^flox/flox^* mice (p53KO) followed by crossing with *Alb-Cre* mice. To generate DKO*^Alb-creERT2^* mice, *p53*KO mice were crossed with transgenic mice carrying *Alb-creERT2* (Schuler et al., 2004), which were a gift from Dr. Czaja (Albert Einstein College of Medicine, NY). For tamoxifen inducible KO experiments, mice received a single intraperitoneal injection of 100 mg/kg tamoxifen. For acute injury experiments by carbon tetrachloride, mice received a single intraperitoneal injection of 1 ml/kg carbon tetrachloride in corn oil or corn oil for control. Mice were sacrificed from 11 am to 1 pm. Mice were maintained under controlled environmental conditions under a 12-h light /dark cycle and allowed ad libitum access to water and standard laboratory diet. Both male and female mice were used for this study. All animal experiments were approved and performed in accordance with the Institutional Animal Care and Use Committee (IACUC) at Rockefeller University.

### Liver function tests

Whole blood from mice was collected via retro-orbital puncture in BD Microtainer blood collection tubes and incubated at room temperature for 30 min, followed by centrifugation at 12,000 x g for 2 min. Triglycerides (TRIG) were measured at the Department of Comparative Pathology in Memorial Sloan Kettering Cancer Center.

### Protein isolation and immunoblot

Liver tissues were homogenized in sucrose A buffer (15 mM Hepes [pH7.9], 60 mM KCl, 2 mM EDTA [pH 8.0], 0.32 mM sucrose) and layered onto sucrose B buffer (15 mM Hepes [pH7.9], 60 mM KCl, 2 mM EDTA [pH 8.0], 30% sucrose), followed by centrifugation at 3,000 rpm for 15 min. For whole cell extracts, the pellet was re-suspended in sucrose A buffer containing 0.2% Triton X-100 and layered onto sucrose B buffer. After centrifugation, the pellet was re-suspended in urea/SDS loading buffer (8M urea, 0.2 M Tris-HCl [pH 6.8], 1 mM EDTA [pH 8.0], 5% SDS, 1.5% DTT, 1% bromophenol blue) and incubated at 40 °C in a Thermomixer R (Eppendorf) for 0.5 h. For nuclear pellet analysis, the pellet was re-suspended in 1 X NUN buffer (1 M urea, 20 mM Hepes [pH7.9], 7.5 mM MgCl_2_, 0.2 mM EDTA [pH 8.0], 300 mM NaCl, 1 % NP40, 1 mM DTT) and incubated with rotation at 4 °C for 5 min, followed by centrifugation at 1,000 x g for 3 min. The pellet was re-suspended in urea/SDS loading buffer and incubated at 40 °C in a Thermomixer R for 0.5 h. For Pol II purification, frozen liver tissues were homogenized in sucrose A buffer and layered onto sucrose B buffer, followed by centrifugation at 3,000 rpm for 15 min. The pellet was re-suspended in 1 pellet volume of 1 x NUN buffer and incubated on ice for 10 min, followed by centrifugation at 13, 000 rpm for 15 min. The supernatant was dialyzed in TGEA buffer (20 mM Tris-HCl [pH 7.9] at 4 °C, 25% glycerol, 0.1 mM EDTA [pH 8.0], 2 mM DTT, 0.5 mM PMSF, 0.1 M ammonium sulfate) and fractionated on a DEAE Sephacel column (GE Healthcare). The flow-through fractions were further fractionated on a phosphocellulose 11 column. For immunoblot, the following antibodies were used: anti-Gdown1 (Jishage et al., 2012), anti-RPB3 (Bethyl Laboratories, A303-771A), anti-cyclinD1 (abcam, ab16663), anti-p21 (abcam, ab188224), anti-phospho STAT3 (Cell Signaling Technology, 9145), anti-NF-κB p65 (Cell Signaling Technology, 8242), anti-β-catenin (abcam, ab32572).

### Immunohistochemistry

Tissues were fixed in 4% paraformaldehyde in phosphate-buffered saline (PBS) for 24 hr, and then processed for embeddedding in paraffin. Tissue sections (5-μm) were cut and deparaffinized in xylene, followed by serial (100%, 95%, 70%, 50%) alcohol washes to rehydrate, and were stained with hematoxylin and eosin (H&E). The rehydrated sections were boiled in antigen unmasking solution (Table 1) (Vector Laboratories) in a pressure cooker for 15 min. After incubation in 0.1% Triton X-100 in PBS for 10 min, primary antibodies in 5% goat serum and 0.1% Triton X-100 in PBS at the indicated dilutions (Table 1) were applied to the sections and were incubated for overnight at 4 °C. The detection was performed using VECTASTAIN Elite ABC HRP kit (Vector Laboratories) according to manufacturer’s instructions and the sections were counterstained with hematoxylin. Tunel assays were performed as described previously (Gavrieli et al., 1992). For Sirius red staining, following deparaffinization, the sections were stained with Picro-Sirius red solution containing 0.1% Direct Red 80 (#365548, Sigma) and 0.1% fast green (F7252, Sigma) in 1.2% saturated aqueous picric acid solution (#197378, Sigma) for 1 hr. Sections were rinsed with water, dehydrated, and mounted with VECTA Mount (Vector Laboratories).

**Table 1.** Primary antibody

| **Antibody** | **Source** | **Identifier** | **Host animal** | **Dilution** | **Antigen retrieval** |
| --- | --- | --- | --- | --- | --- |
| Ki67 | abcam | ab16667 | Rabbit | 1:100 | pH 6.0 |
| CyclinD1 | Thermo | RM-9104-S1 | Rabbit | 1:100 | pH 6.0 |
| SMA | abcam | ab32575 | Rabbit | 1:500 | pH 6.0 |
| KRT19 | DSHB | Troma-III | Rat | 1:200 | pH 9.0 |
| Phospho-H3 | abcam | ab5176 | Rabbit | 1:2000 | pH 6.0 |
| Glutamine synthetase | Sigma | G2781 | Rabbit | 1:40,000 | pH 6.0 |
| p21 | abcam | ab188224 | Rabbit | 1:1000 | pH 6.0 |
| p53 | Leica | NCL-L-P53-CM5p | Rabbit | 1:800 | pH 6.0 |
| CD44 | Cell Signaling Technology | 37259 | Rabbit | 1:500 | pH6.0 |
| Phospho-c-Jun | Cell Signaling Technology | 3270 | Rabbit | 1:200 | pH6.0 |
| AFP | Biocare Medical | CP028A | Rabbit | 1:100 | pH6.0 |

### RNA isolation and RT-PCR

Liver tissue samples were homogenized in 0.7 ml of TRIZOL (ThermoFisher). RNA was purified by RNA Clean & concentrator (Zymo research). RNA was reverse-transcribed using Superscript III First Strand Synthesis kit (Invitrogen). Real-time PCR was performed on an Applied Biosystems 7300 Real Time PCR System using QuantiTect SYBER Green mix. Changes in mRNA expression were calculated using the ΔΔ Ct method and are presented as fold-change in relation to expression of the *Actb* gene.

### Microarray and RNA-seq analysis

RNA quality was assessed using an Agilent Bioanalyzer. Microarray analysis was performed using Affymetrix Gene Chip Gene 2.0 ST arrays according to the manufacturer’s instructions. The data expressed as CEL files were normalized by the robust multiarray average method with the Expression Console software (Affymetrix). For RNA-seq, an RNA library was generated using SMARTer® Stranded Total RNA Sample Prep Kits (Takara Bio) according to the manufacturer’s instructions. Library quality was evaluated using the Bioanalyzer and sequenced on Illumina NextSeq High (75 bp single-end). Reads were processed using Trimmomatic (version 0.39) (Bolger et al., 2014) and aligned to the mouse genome (mm10) using hisat2 (version 2.1.0) (Kim et al., 2015). Aligned reads were counted using featureCounts (Subread version 1.6.4) (Liao et al., 2014), and differential expression analysis was performed using DESeq2 (version 1.22.2) (Love et al., 2014). GO pathway analysis was performed by DAVID Bioinformatics Resources 6.8, and gene set enrichment analysis was performed by GSEA (Subramanian et al., 2005).

### ChIP-seq analysis

Liver tissues were homogenized in sucrose A buffer (15 mM Hepes [pH7.9], 60 mM KCl, 2 mM EDTA, 0.32 mM sucrose) and layered onto sucrose B buffer (15 mM Hepes [pH7.9], 60 mM KCl, 2 mM EDTA, 30% sucrose), followed by centrifugation at 3,000 rpm for 15 min. The pellet was re-suspended in sucrose A buffer containing 0.2% Triton X-100 and layered onto sucrose B buffer. After centrifugation, the pellet was re-suspended in MNase buffer and incubated with MNase (New England Biolabs) for 10 min at 37 °C. The MNase-digested chromatin was further sonicated briefly and centrifuged at 14,000 rpm for 15 min. The supernatant was diluted in Buffer C (20 mM Hepes [pH7.9], 100 mM KCl, 2 mM EDTA, 20% glycerol, 0.1% NP40) and pre-cleared with Dynabeads Protein A for 2 hours at 4 °C. Anti-Gdown1 antibodies (Jishage et al., 2012) or anti-RPB3 antibodies (Bethyl Laboratories, A303-771A) that were pre-incubated with Dynabeads Protein A, were added to the pre-cleared chromatin, followed by rotation overnight at 4 °C. The Dynabeads were washed three times in Buffer C for 10 min three times and rinsed in TE buffer (10 mM Tris [pH8], 1 mM EDTA), followed by two elutions in TE buffer containing 1% SDS at 65 °C for 15 min twice. The combined eluates were treated with RNase A (10 μg/ml) followed by proteinase K (0.1 mg/ml), and DNA was extracted by QIAquick PCR purification kits (Qiagen). DNA quality was assessed using an Agilent Bioanalyzer, and quantified using a Qubit Fluorometer (Invitrogen). ChIP-seq libraries were generated using DNA SMART ChIP-Seq kits (Takara Bio) according to the manufacturer’s instructions. The library quality was evaluated using the Bioanalyzer and sequenced on Illumina NextSeq High (75 bp single-end). Reads were trimmed using Trimmomatic (version 0.39) (Bolger et al., 2014) and aligned to the mouse genome (mm10) using Bowtie2 (version 2.3.5) (Langmead and Salzberg, 2012), followed by process using Samtools (Li et al., 2009). Peak calling was performed using MACS2 (Zhang et al., 2008) for narrow peaks and HOMER (Heinz et al., 2010) for broad peaks, and peak annotation was performed using HOMER. Aligned reads were normalized using deepTools (Ramirez et al., 2016) and visualized in the WashU Epigenome Browser (Zhou and Wang, 2012).

### Quantification and statistical analysis

Unpaired two-tailed *t* test was used to analyze differences between two groups. P-values less than 0.05 were considered statistically significant. The analysis was performed using GraphPad Prism8 (GraphPad software).

## Acknowledgements

This study was supported by National Institutes of Health grants R01 CA202245 and R01 CA129325 to R.G.R. We thank Sohail Malik for discussions and critical reading of the manuscript.

## Author Contributions

M.J. performed analyses for KO*^Alb-cre^*, DKO*^Alb-cre^*, and DKO*^Alb-creERT2^* liver by histology, biochemical analysis, and RNA expression analysis. K.I. assisted KO*^Alb-cr^*^e^ liver analysis. C.C. performed the ChIP-seq analysis. X.Y. assisted the early embryonic analysis. M.Y. designed and performed the screening of Gdown1 KO mESC. M.J. and R.G.R. wrote the manuscript with input from all co-authors.

## Additional Resources

All raw data are deposited in GEO and the accession number is GSE144212.

