## Supplemental information for "Transcriptional down-regulation of metabolic genes by Gdown1 ablation induces quiescent cell re-entry into the cell cycle"

**A**

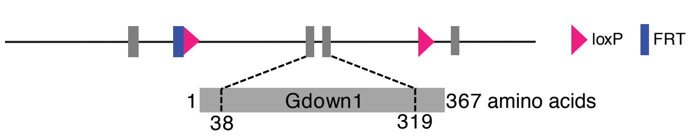


**B**

| | Gdown1 $^{f/f}$ | Gdown1 $^{f/-}$ | Gdown1 $^{-/-}$ | Total |
|---|---|---|---|---|
| E2.5 | 8 | 38 | 9 | 55 |
| E3.5 | 12 | 31 | 4 | 47 |
| E10.5 | 11 | 40 | 0 | 42 |
| P21 | 25 | 42 | 0 | 67 |

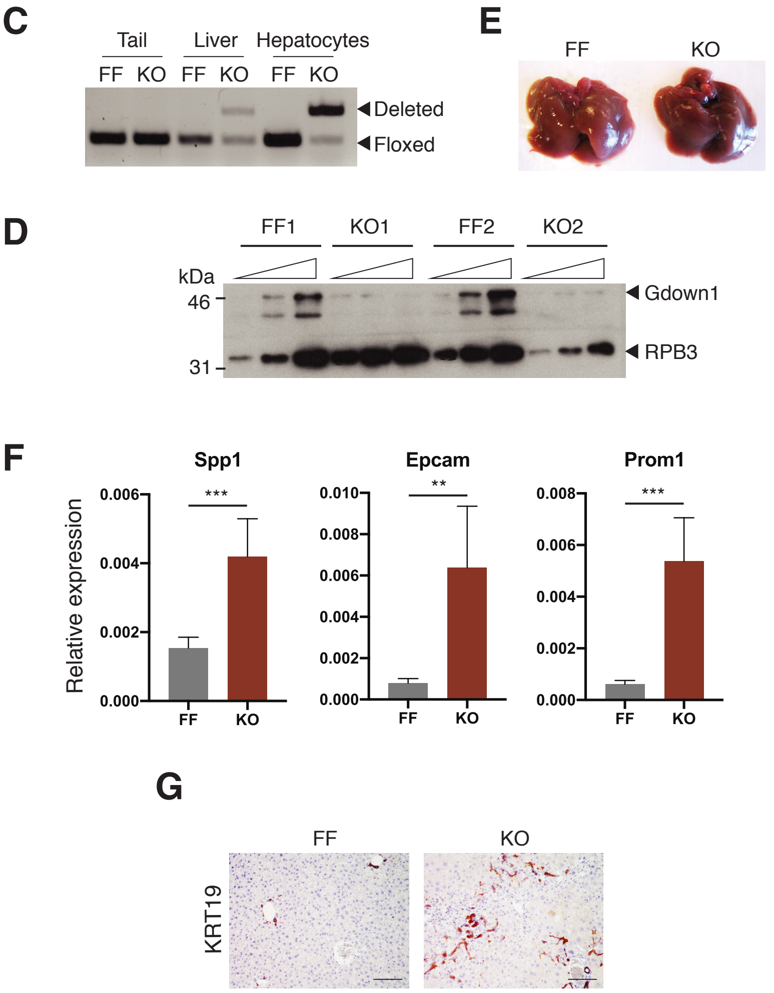


**Supplementary Figure 1. Related to Figure 1. Loss of Gdown1 activates p53 signaling pathway.** (A) Schematic representing *Gdown1* targeting construct. (B) The numbers of embryos at embryonic day 2.5 (E2.5) to E10.5 and offspring at postnatal day 21 from intercrossing of *Gdown1*$^{flox/-}$ mice. (C) Genotyping analysis by PCR for detecting *Gdown1* deleted and floxed allele in the indicated tissues from *Gdown1*$^{flox/flox}$ (FF) and *Gdown1*$^{flox/flox;Alb\text{-}cre}$ (KO) mice. (D) Gdown1 expression in FF and KO liver at 8 weeks of age. Liver nuclear pellets were analyzed by immunoblot. RPB3 serves as a loading control. (E) Livers from FF and KO mice. (F) Relative mRNA expression of the indicated genes analyzed by Real-time qPCR. Data are presented with mean and SD (n = 5 mice at 6W per group). ***$P < 0.001$, ****$P < 0.0001$ in unpaired two-tailed t-test. (G) Representative liver histology for KRT19-positive hepatocytes in FF and KO mice at 8W.

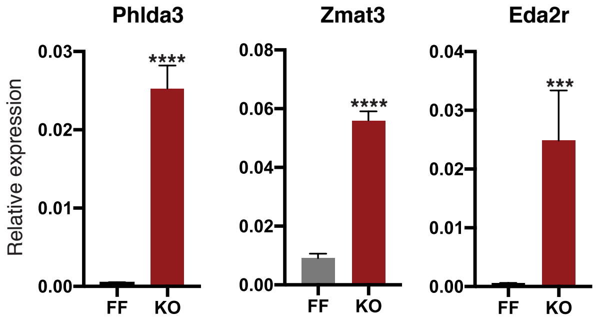


**Supplementary Figure 2. Related to Figure 2. Gdown1 KO hepatocytes re-enter into the cell cycle.**
Relative mRNA expression of p53 target genes analyzed by Real-time qPCR. Data are presented with mean and SD (n = 4-5 mice at the ages of 6W per group). ***$P$ < 0.001, ****$P$ < 0.0001 in unpaired two-tailed t-test.

**A**

**Up-regulated genes**

| Term | Count | % | PValue |
|---|---|---|---|
| GO:0007049~cell cycle | 159 | 8.7 | 1.76E-36 |
| GO:0007067~mitotic nuclear division | 90 | 4.9 | 2.19E-28 |
| GO:0007155~cell adhesion | 120 | 6.5 | 1.19E-25 |
| GO:0051301~cell division | 101 | 5.5 | 9.67E-25 |
| GO:0002376~immune system process | 90 | 4.9 | 7.29E-18 |
| GO:0006260~DNA replication | 41 | 2.2 | 1.23E-13 |
| GO:0016477~cell migration | 52 | 2.8 | 3.73E-13 |
| GO:0007159~leukocyte cell-cell adhesion | 17 | 0.9 | 4.92E-12 |
| GO:0007059~chromosome segregation | 31 | 1.7 | 5.10E-11 |

**Down-regulated genes**

| Term | Count | % | PValue |
|---|---|---|---|
| GO:0055114~oxidation-reduction process | 146 | 12.2 | 2.79E-48 |
| GO:0006629~lipid metabolic process | 103 | 8.6 | 2.79E-35 |
| GO:0008152~metabolic process | 98 | 8.2 | 2.24E-31 |
| GO:0008202~steroid metabolic process | 37 | 3.1 | 1.13E-23 |
| GO:0008203~cholesterol metabolic process | 37 | 3.1 | 1.25E-22 |
| GO:0006631~fatty acid metabolic process | 42 | 3.5 | 1.32E-17 |
| GO:0009813~flavonoid biosynthetic process | 14 | 1.2 | 5.03E-12 |

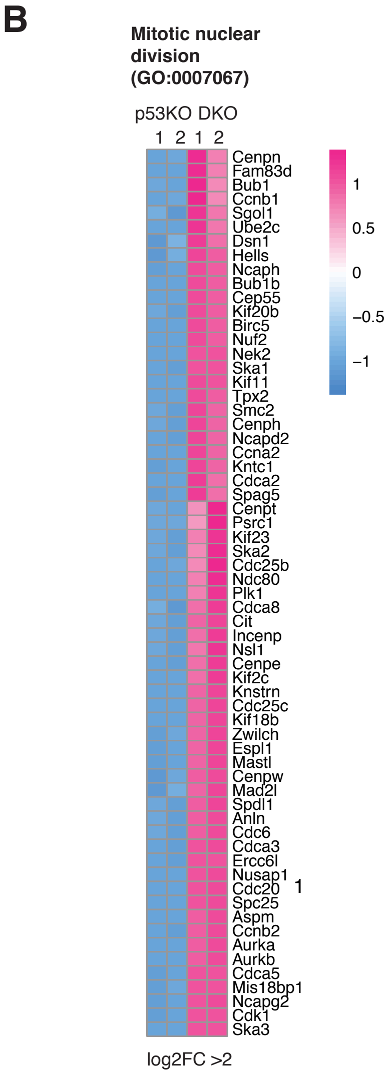


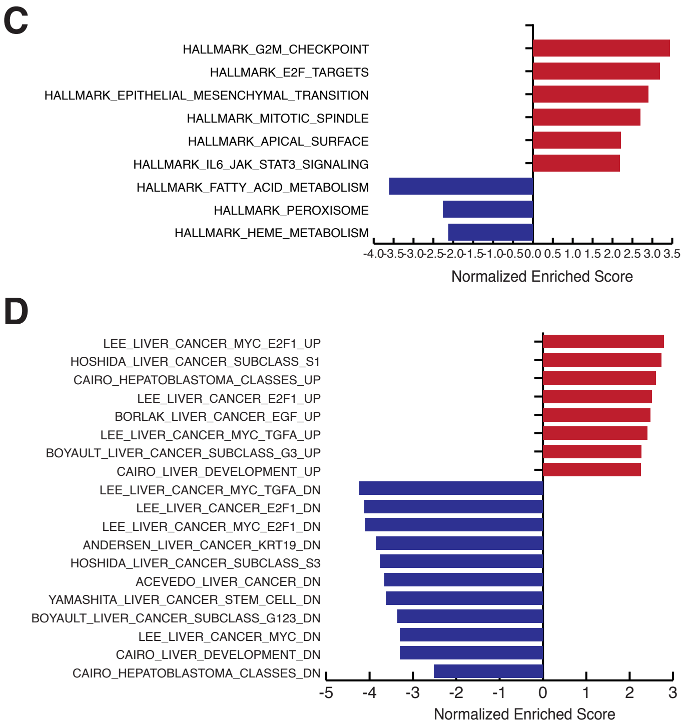


**Supplementary Figure 3. Related to Figure 3. Gdown1 KO causes dysregulated cell cycle progression in the absence of p53.**

(A) Gene ontology analysis for differentially expressed genes in DKO liver. (B) A heat map for mitotic nuclear division genes that were up-regulated in DKO liver. (C) (D) Normalized enrichment scores for DKO liver relative to p53 KO liver compared with hallmark gene sets (C) or human HCC gene sets (D) by GSEA.

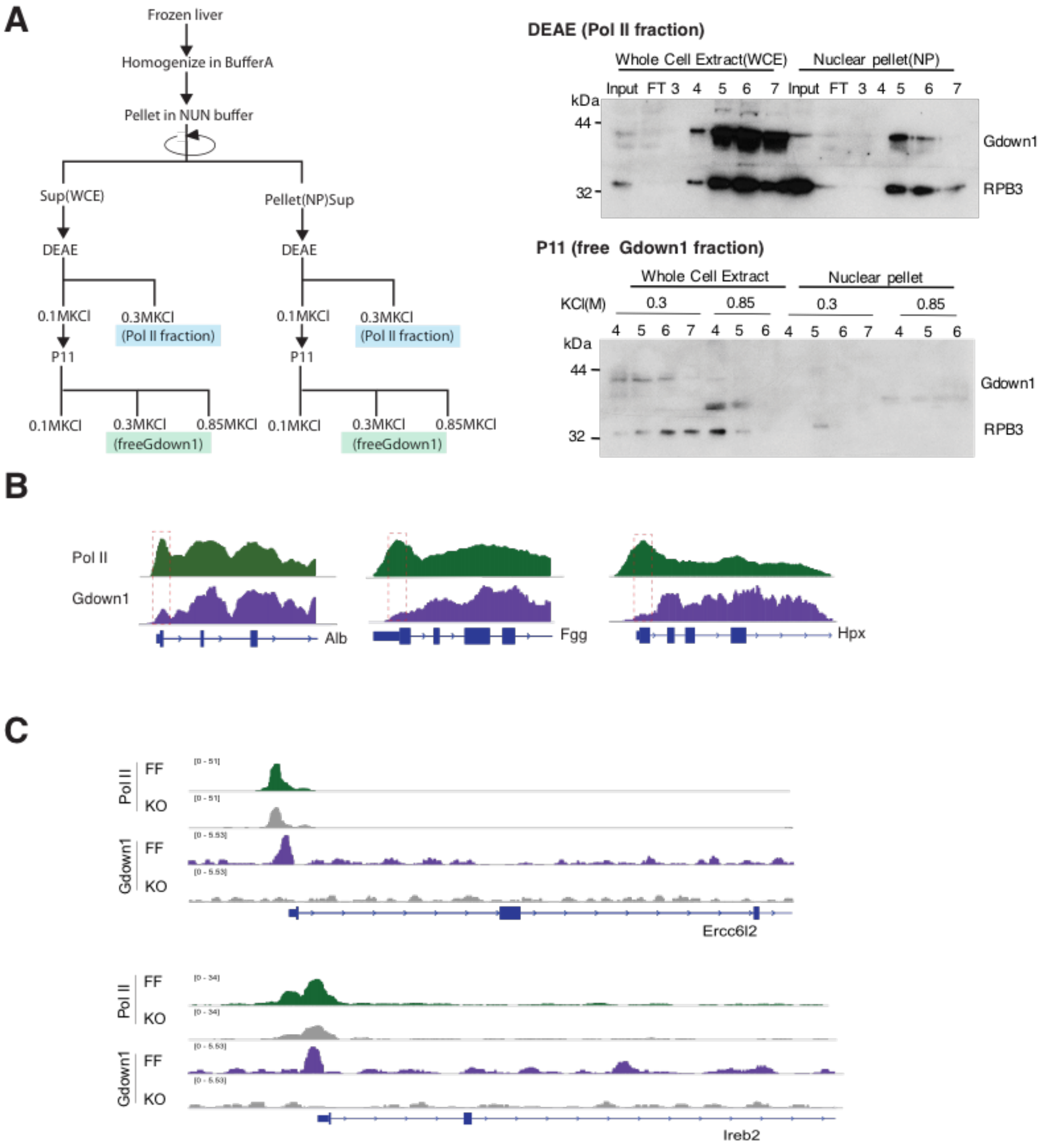


**Supplementary Figure 4. Related to Figure 4. Gdown1 is associated with elongating Pol II on genes that actively transcribed in the liver.**
(A) Left: Schematic of Pol II(G) purification. Right: Immunoblot for Pol II(G) fractions analyzed by ion-exchange chromatography. The liver whole cell extracts or nuclear pellet were subjected to anion-exchange chromatography (DEAE) and the flow through fractions were further analyzed by cation-exchange chromatography (P11). (B) ChIP-seq profiles generated with Gdown1 and RPB3 (Pol II) antibodies at the promoter proximal regions of indicated genes in FF. (C) ChIP-seq profiles generated with Gdown1 and RPB3 (Pol II) antibodies at the promoter regions of indicated genes in FF and KO liver.

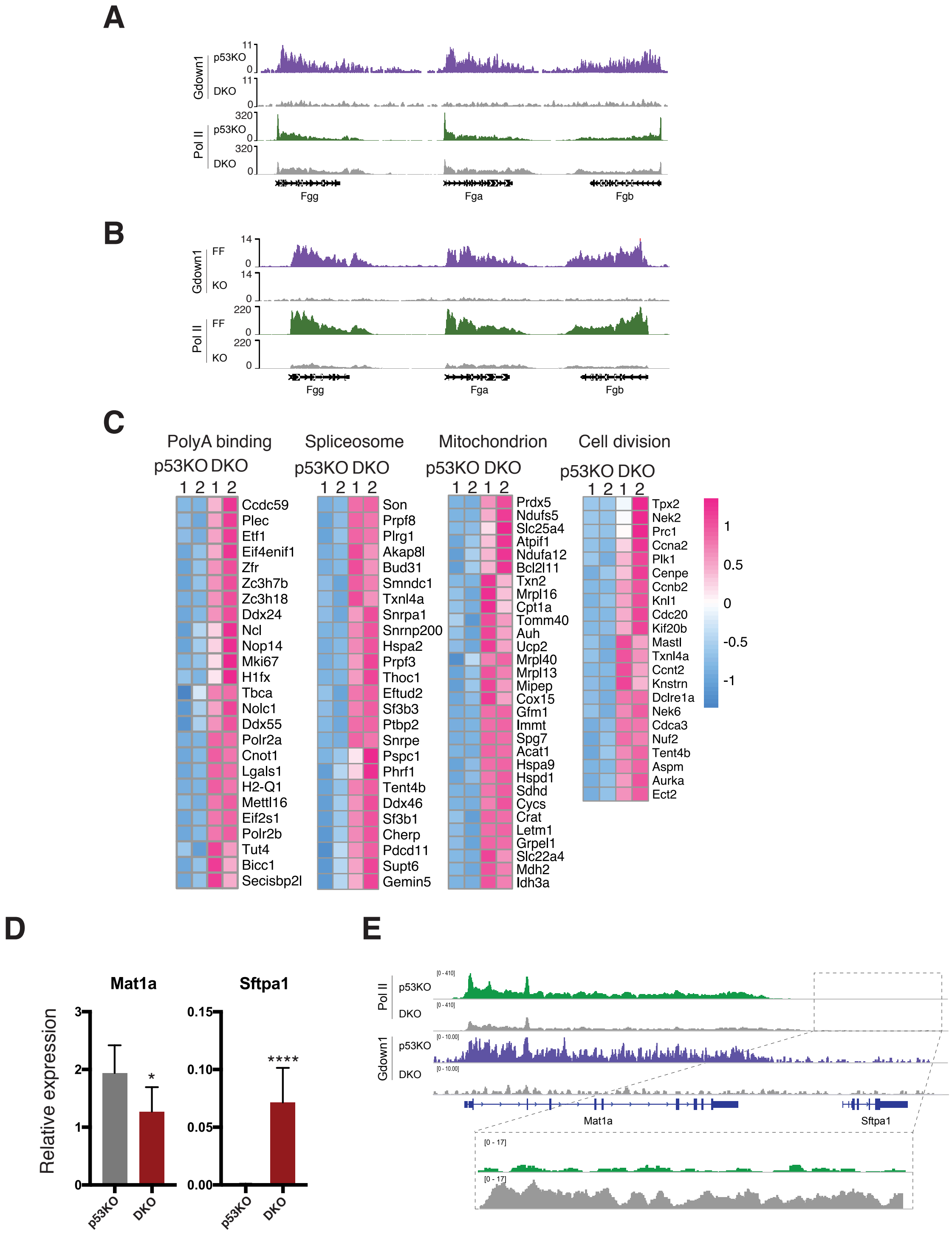
A
Gdown1
p53KO
DKO
Pol II
p53KO
DKO
11
0
320
Fgg
Fga
Fgb
B
Gdown1
FF
KO
Pol II
FF
KO
14
0
220
Fgg
Fga
Fgb
C
PolyA binding
p53KO DKO
1 2 1 2
Ccdc59
Plec
Etf1
Eif4enif1
Zfr
Zc3h7b
Zc3h18
Ddx24
Ncl
Nop14
Mki67
H1fx
Tbca
Nolc1
Ddx55
Polr2a
Cnot1
Lgals1
H2-Q1
Mettl16
Eif2s1
Polr2b
Tut4
Bicc1
Secisbp2l
Spliceosome
p53KO DKO
1 2 1 2
Son
Prpf8
Plrg1
Akap8l
Bud31
Smndc1
Txnl4a
Snrpa1
Snrnp200
Hspa2
Prpf3
Thoc1
Eftud2
Sf3b3
Ptbp2
Snrpe
Pspc1
Phrf1
Tent4b
Ddx46
Sf3b1
Cherp
Pdcd11
Supt6
Gemin5
Mitochondrion
p53KO DKO
1 2 1 2
Prdx5
Ndufs5
Slc25a4
Atpif1
Ndufa12
Bcl2l11
Txn2
Mrpl16
Cpt1a
Tomm40
Auh
Ucp2
Mrpl40
Mrpl13
Mipep
Cox15
Gfm1
Immt
Spg7
Acat1
Hspa9
Hspd1
Sdhd
Cycs
Crat
Letm1
Grpel1
Slc22a4
Mdh2
Idh3a
Cell division
p53KO DKO
1 2 1 2
Tpx2
Nek2
Prc1
Ccna2
Plk1
Cenpe
Ccnb2
Knl1
Cdc20
Kif20b
Mastl
Txnl4a
Ccnt2
Knstrn
Dclre1a
Nek6
Cdca3
Nuf2
Tent4b
Aspm
Aurka
Ect2
1
0.5
0
-0.5
-1
D
Mat1a
Sftpa1
Relative expression
0
1
2
3
0.00
0.05
0.10
0.15
*
****
p53KO
DKO
E
Pol II
p53KO
DKO
Gdown1
p53KO
DKO
Mat1a
Sftpa1

**Supplementary Figure 5. Related to Figure 5. Expression of Gdown1 target genes is inversely correlated to expression of cell cycle-involved genes.**
(A) ChIP-seq profiles generated with Gdown1 and RPB3 (Pol II) antibodies at the indicated genes in p53KO or DKO$^{Alb\text{-}creERT2}$ liver at 24 hours after tamoxifen injection. (B) ChIP-seq profiles generated with Gdown1 and RPB3 (Pol II) antibodies at the indicated genes in FF or KO$^{Alb\text{-}cre}$ liver at 7W. (C) Heat maps for the up-regulated genes in DKO$^{Alb\text{-}creERT2}$ liver that were grouped by GO analysis. (D) Relative mRNA expression of the indicated genes in p53KO or DKO$^{Alb\text{-}creERT2}$ liver at 24 hours after tamoxifen injection analyzed by Real-time qPCR. Data are presented with mean and SD (n = 3-7 mice per group). *$P$ < 0.05, ****$P$ < 0.0001 in unpaired two-tailed t-test. (E) ChIP-seq profiles generated with Gdown1 and RPB3 (Pol II) antibodies at the indicated genes in p53KO or DKO$^{Alb\text{-}creERT2}$ liver at 24 hours after tamoxifen injection.

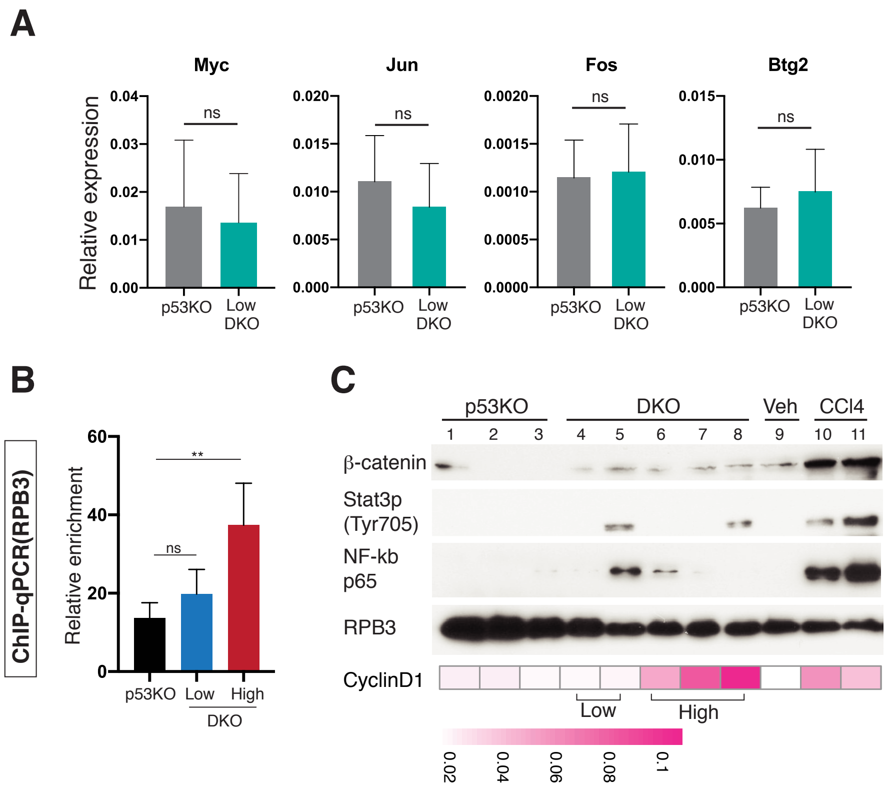


**Supplementary Figure 6. Related to Figure 6. Down-regulation of highly expressed genes in the liver contributes to hepatocytes re-entry into the cell cycle**

(A) Relative mRNA expression of the indicated genes analyzed by Real-time qPCR. Control, p53KO treated with tamoxifen for 16 hours (n = 7). Low, DKO$^{Alb\text{-}creERT2}$ treated with tamoxifen for 16 hours expressed at the low level of cyclin D1 (n = 6). ns, not significant. (B) Relative enrichment of Pol II at the promoter region for cyclin D1 analyzed by Real-time qPCR. Control, p53KO mice treated with tamoxifen for 16 hours (n = 4 mice per group). Low or High, DKO$^{Alb\text{-}creERT2}$ mice treated with tamoxifen for 16 hours and expressing low or high levels of cyclin D1 (n = 6 or 4 mice per group). (C) Nuclear expression of the indicated proteins in livers from p53KO (lanes 1-3) or DKO$^{Alb\text{-}creERT2}$ mice treated with tamoxifen for 16 hours (lane 4-8) or from FF mice treated either with corn oil (lane 9) or with $CCl_4$ for 12 hours (lane 10 and 11). Liver nuclear pellets were analyzed by immunoblot. RPB3 was used for normalizing. The expression levels of cyclin D1 mRNA are presented in a heat map.

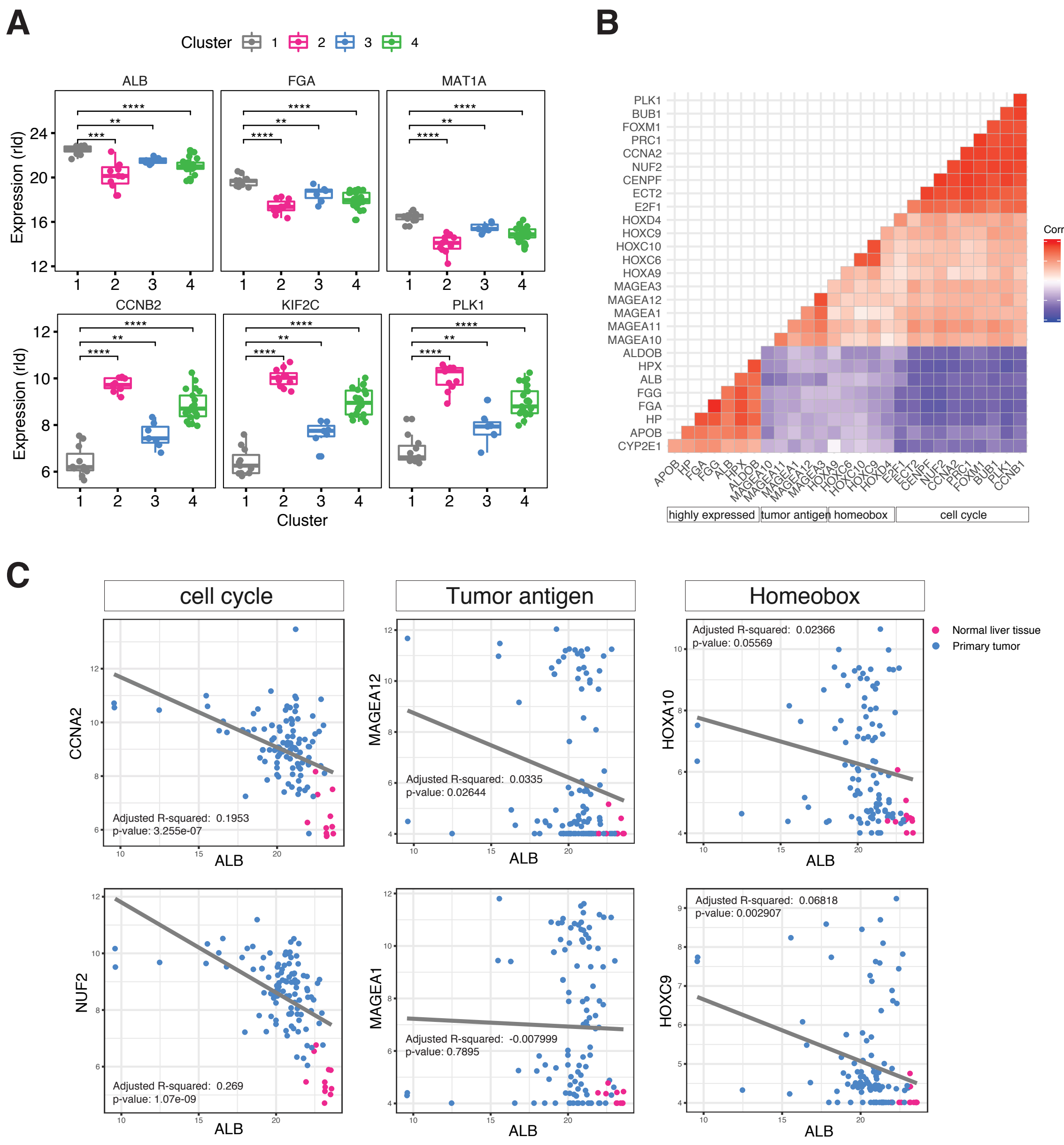


**Supplementary Figure 7. Related to Figure 7. Expression of metabolic genes is inversely correlated to expression of cell cycle-involved genes in HCC**

(A) Comparison of the indicated gene expression by clusters that were analyzed in Figure 7A. (B) Correlation plot with the expression of the indicated genes that are grouped by function. (C) Scatter plot and regression line with expressions of ALB and the indicated genes in normal liver tissue (shown in pink) and HCC cohorts with p53 mutations (shown in blue).
